# Landscape of Evolutionary Arms Races between Transposable Elements and KRAB-ZFP Family

**DOI:** 10.1101/2024.05.20.595046

**Authors:** Masato Kosuge, Jumpei Ito, Michiaki Hamada

## Abstract

Transposable elements (TEs) are mobile parasitic sequences that have expanded within the host genome. It has been hypothesized that host organisms have expanded the Krüppel-associated box-containing zinc finger proteins (KRAB-ZFPs), which epigenetically suppress TEs, to counteract disorderly TE transpositions. This process is referred to as the evolutionary arms race. However, the extent to which this evolutionary arms race occurred across various TE families remains unclear. In the present study, we systematically explored the evolutionary arms race between TE families and KRAB-ZFPs using public ChIP-seq data. We discovered and characterized new instances of evolutionary arms races with KRAB-ZFPs in endogenous retroviruses. Furthermore, we found that the regulatory landscape shaped by this arms race contributed to the gene regulatory network. In summary, our results provide insight into the impact of the evolutionary arms race on TE families, the KRAB-ZFP family, and host gene regulatory networks.

## Introduction

Transposable elements (TEs) are mobile parasitic genetic sequences that comprise approximately 46% of the human genome^1^. Although most insertions of TEs near genes are likely harmful or neutral to their host organisms, TEs have significantly influenced the evolution of their host organisms through transpositions^1^. TEs possess numerous binding sites for transcription factors (TFs) and their insertion generates new binding sites for TFs near genes^2,3^. Some new TE insertions can function as cis-regulatory elements, such as enhancers or alternative promoters of genes near their insertion sites, thereby altering the expression patterns of these genes^4,5^. Furthermore, TEs acquire or lose the binding sites of specific TFs during evolution, resulting in each TE subfamily having distinct expression profiles and effects on nearby genes^2,6^. Therefore, uncovering the evolution of TEs is crucial for understanding the evolution of host organisms.

Krüppel-associated box-containing zinc finger proteins (KRAB-ZFPs) are transcriptional repressors that epigenetically suppress the transcription of TEs^7^. The human genome contains approximately 400 genes encoding KRAB-ZFPs ^8–10^. Each KRAB-ZFP comprises one or two KRAB domains that interact with TRIM28 and several zinc finger (ZF) domains that recognize sequences in TEs^7^. KRAB-ZFPs recruit SETDB1, HP1, and the nucleosome remodelling deacetylase complex via TRIM28, which induces H3K9me3 and DNA methylation, thereby epigenetically suppress TEs^11–13^. In humans, depletion of KRAB-ZFP and TRIM28 leads to de-repression of TEs in developmental stages^14,15^. In addition, in mice, deletion of the KRAB-ZFP cluster slightly induced transpositions of ERVs^16^. Thus, KRAB-ZFPs play a crucial role in the suppression of TEs.

It has been hypothesized that the significant expansion of KRAB-ZFPs is the result of an evolutionary arms race with TEs^7^. This arms race is a co-evolutionary process among completers. In the context of KRAB-ZFPs and TEs, the proposed evolutionary scenario is as follows. As new TEs emerge and proliferate within the host genome, KRAB-ZFPs that specifically suppress these TEs emerge through gene duplication. Subsequently, TEs acquire mutations in the binding sites of KRAB-ZFPs, enabling them to avoid suppression by KRAB-ZFPs. This evolutionary scenario is supported by reports indicating that TEs are often targeted by KRAB-ZFPs that emerged concurrently^8,9,17^. Additionally, the long interspersed nuclear element (LINE) 1 family evades suppression by a specific KRAB-ZFP, ZNF93, through deletions at its binding sites^18^. However, the extent to which this evolutionary arms race has occurred in other TE families remains unclear.

Such a relationship between TEs and KRAB-ZFPs has been proposed to be co-opted as part of the gene regulatory network^8,19^. KRAB-ZFPs regulate gene expression by suppressing TEs that function as cis-regulatory elements^8,20-^22. Perturbation of KRAB-ZFP or TRIM28 expression led to aberrant expression of nearby genes^23–25^. However, the effect of the evolutionary arms race between TE families and KRAB-ZFPs on the gene regulatory network is still not well understood.

In this study, we aimed to comprehensively investigate the evolutionary arms race between TE families and KRAB-ZFPs using multi-layered analyses: (1) identification of KRAB-ZFP targets by leveraging publicly available ChIP-seq data, (2) comparison of the evolutionary ages between TE subfamilies and KRAB-ZFPs, and (3) phylogenetic analyses of the TE family (**Fig. 1**). Accordingly, we reconstructed and characterized the evolutionary arms race with KRAB-ZFPs in several TE families, which has not been previously reported. Finally, we provide evidence that these arms race dynamics have potentially influence the evolution of the host gene regulatory network. Our findings illustrate the co-evolutionary relationship between TE and KRAB-ZFPs, offering fascinating insights into TE-host interactions.

**Fig. 1.**
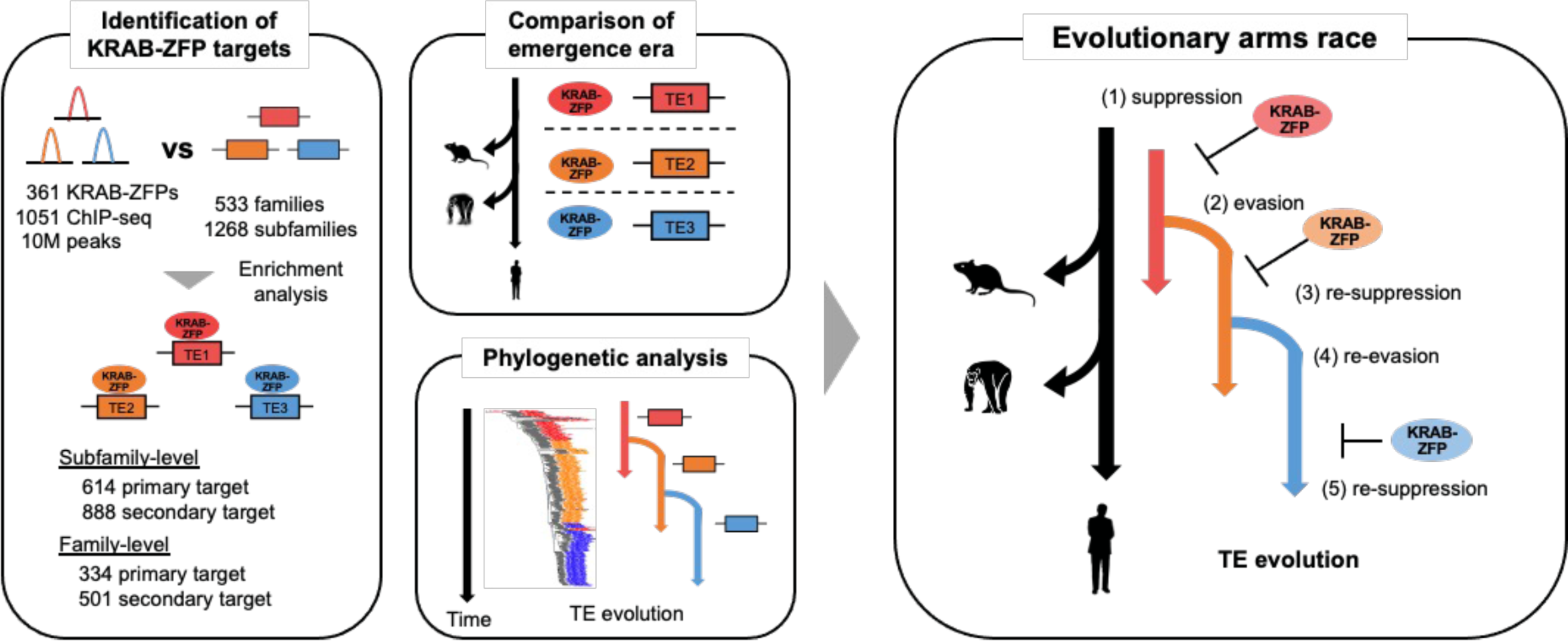
Overview of the study design and the evolutionary arms race model between TE family and KRAB-ZFPs. Left panel: Schematic of the main analyses performed in this study, including identification of KRAB-ZFP targets, comparison of the evolutionary age between KRAB-ZFPs and TE families, and phylogenetic analysis of TE families. Right panel: The evolutionary arms race between the TE family and KRAB-ZFPs. This arms race involves recurrent cycles of (1) TE suppression by KRAB-ZFPs, (2) TE evasion from KRAB-ZFP suppression, (3) re-suppression by existing or newly emerged KRAB-ZFPs, (4) further TE evasion, and (5) re-suppression by additional KRAB-ZFPs.

## Results

### Characterization of the relationship between TE families and KRAB-ZFPs

To comprehensively characterize the relationship between TE families and KRAB-ZFPs, we first collected and processed a large dataset of KRAB-ZFP ChIP-seq experiments encompassing 361 KRAB-ZFPs from 1,051 samples (**Supplementary Fig. 1a, b** and **Supplementary Table 1**). We obtained consensus sequences and metadata for 1170 TE subfamilies belonging to 533 TE families from Dfam^26^. Given the association between alterations in TE sequences over evolution and KRAB-ZFP binding, we focused on the differences in sequences and KRAB-ZFP binding among the TE subfamilies^8,18,27^. To comprehensively explore the evolutionary arms race with KRAB-ZFPs, we developed a novel subfamily classification pipeline based on the genetic distance between TE copies and performed subfamily classification in TE families that previously lacked subfamily classification (344 families, 65 %) (**Supplementary Fig. 2, 3** and **Methods**). By applying our pipeline to the TE families, we successfully divided the 59 families that met our inclusion criteria into subfamilies (**Supplementary Fig. 2c, d**). The final dataset comprised 533 TE families and 1,268 subfamilies (**Supplementary Data 1**).

We conducted enrichment analyses to examine the TE targets of KRAB-ZFPs and identified 614 primary and 888 secondary targets of KRAB-ZFPs (**Supplementary Data 2** and **Methods**). Among 378 KRAB-ZFPs, 266 (74%) targeted at least one subfamily. Additionally, 471 (37%) subfamilies and 146 (27%) families were targeted by at least one KRAB-ZFP, respectively (**Supplementary Fig. 4a, b**). Consistent with previous studies^9^, LINEs and endogenous retroviruses (ERVs) were enriched as targets of KRAB-ZFPs (**Fig.2a**).

Previous studies have described the co-emergence of KRAB-ZFPs and their target TE subfamilies in the same era^8,9,17^. Hence, we obtained the evolutionary ages of the KRAB-ZFPs from two previous studies and inferred the evolutionary ages of their target TE subfamilies (**Supplementary Data 1, 3**). Both the KRAB-ZFPs and TE subfamilies primarily emerged during the evolutionary period of the common ancestors of Eutherians around 105 million years ago (MYA), Simiiformes around 43 MYA, and Catarrhines around 29 MYA. (**Supplementary Fig. 4c**). Consistent with previous studies, the evolutionary ages of KRAB-ZFPs tended to coincide with those of their target TE subfamilies, suggesting that KRAB-ZFPs emerged in response to the emergence and expansion of new TE subfamilies (**Fig. 2b** and **Supplementary Fig. 4d, e**). Unexpectedly, the KRAB-ZFPs that emerged in a common ancestor with non-primates targeted primate-specific TE subfamilies, whereas the reverse pattern was rarely observed. (**Fig. 2b** and **Supplementary Fig. 4f**). These observations imply that KRAB-ZFPs that emerged from the common ancestor with non-primates specifically target and suppress newly emerged TE subfamilies. In summary, these findings support the hypothesis that the KRAB-ZFP family has co-evolved with TEs.

**Fig. 2.**
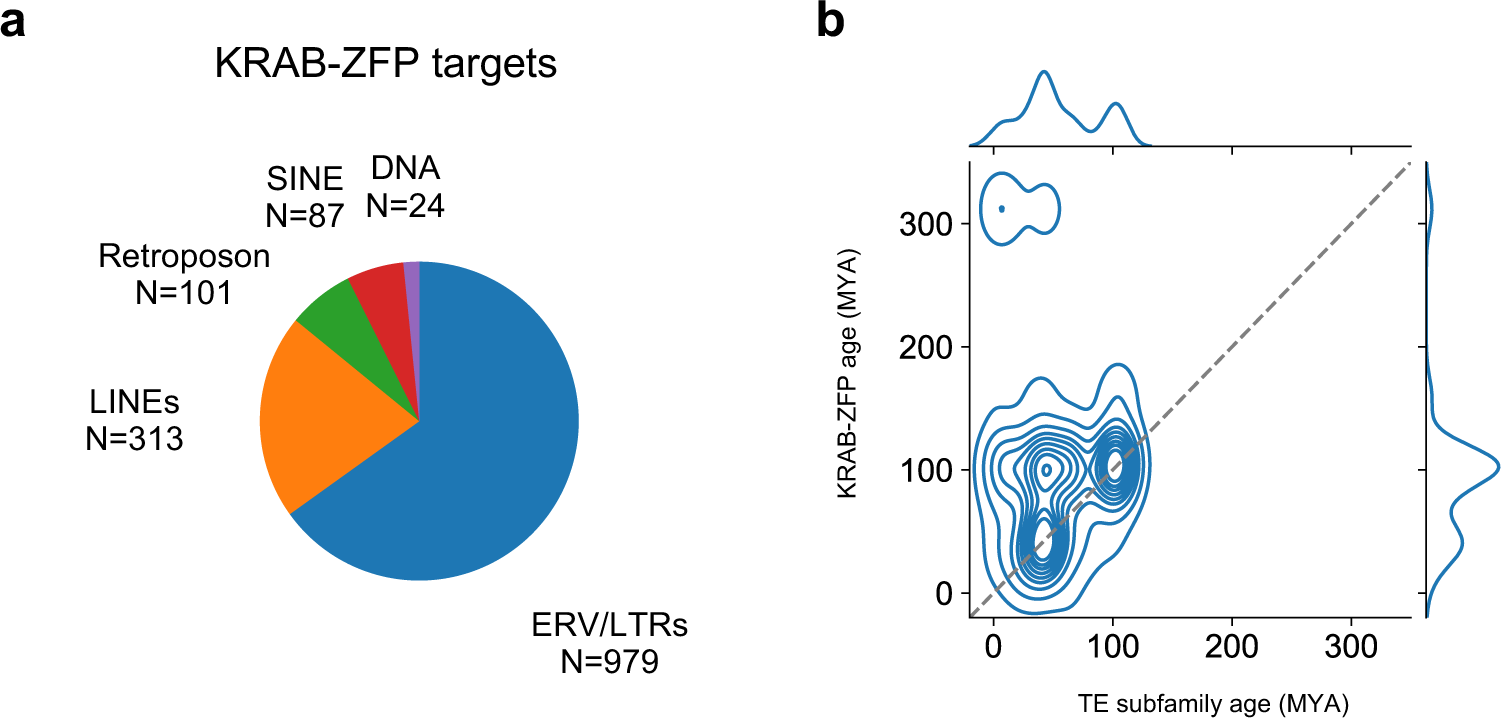
Co-emergence of TE subfamilies and KRAB-ZFPs. **b,** Proportion of TE classes in the identified KRAB-ZFP targets. Pie chart shows the distribution of TE classes, including ERVs (blue), LINEs (orange), SINEs (green), retroposons (red), and DNA transposons (purple). The number of TE subfamilies in each class is indicated by “N=”. b Comparison of evolutionary age (in million years ago, MYA) between the TE subfamily (X-axis) and KRAB-ZFPs (Y-axis) for 614 primary KRAB-ZFP-TE subfamily associations.

### Several TE families have undergone evasion from KRAB-ZFPs

To investigate the evasion of KRAB-ZFP suppression by TE families, we developed a computational screening approach based on the differential binding of KRAB-ZFPs to young and old subfamilies within each TE family (**Fig. 3a**). Our rationale was that if a TE family evolved to evade KRAB-ZFP repression, we would expect to see a significant reduction in KRAB-ZFP binding to its younger subfamilies compared to older ones. Applying this approach to our dataset, we identified 62 evasion candidates that showed reduced binding in younger subfamilies (**Fig. 3b** and **Supplementary Data 4**). As positive controls, loss of binding by ZNF765, ZNF649, and ZNF93 at the L1P_5end was also detected, which is consistent with previous studies (**Supplementary Fig. 5a, b**)^8,18,28^. In addition to the previously reported L1P_5end, evasion candidates were detected in several ERV families and SVAs, suggesting that the evasion of TE families from KRAB-ZFPs occurred across a broad range of TE families.

**Fig. 3.**
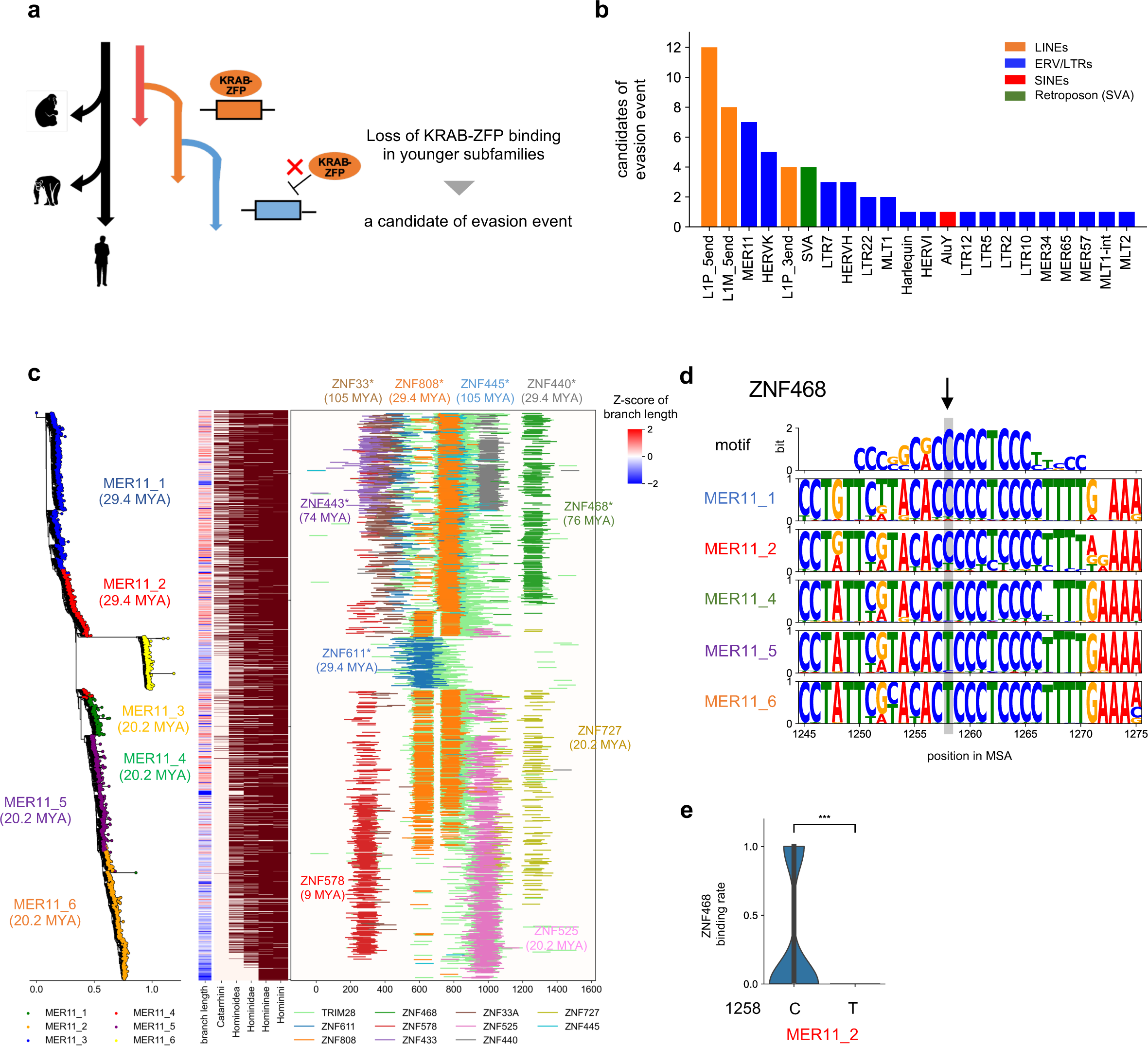
Screening for TE evasion from KRAB-ZFPs and the evolutionary arms race in MER11. **a**, Schematic of the screening approach for identifying potential TE evasion events from KRAB-ZFP repression. Screening was based on the loss of KRAB-ZFP binding in younger TE subfamilies compared with older subfamilies within the same family. **b,** Number of candidate evasion events identified in each TE family. The bar plot shows the number of KRAB-ZFPs that each TE family potentially evaded in our screening approach, including ERVs (blue), LINEs (orange), SINEs (green), and retroposons (red). **c,** Evolutionary arms race in MER11 with 10 KRAB-ZFPs. 7 KRAB-ZFPs potentially evaded by the MER11 family and 3 KRAB-ZFPs primarily targeted the MER11 family. * after gene symbol of KRAB-ZFPs indicates the significance of screening for evasion events. The phylogenetic tree (left) indicates the phylogenetic relationships between the MER11 subfamilies. The ages listed under subfamily names indicate the evolutionary ages of the MER11 subfamilies. The heatmap of the branch length and liftover in the center indicates the insertion date of the MER11 copies. The upper side of the y-axis indicates the older MER11 copies and subfamilies. The plot on the right indicates the positions of TRIM28 and 10 KRAB-ZFP peaks for each MER11 copy. Peaks are depicted in different colors for each KRAB-ZFP gene. **d,** Evasion of ZNF468 due to point mutations. Sequence logos indicate the de novo motif of ZNF468 (top) and the sequence of ZNF468 binding site in the MER11 subfamily. Letters indicate the proportion of bases in each position. Arrow indicates the position potentially associated with the loss of ZNF468 binding. **e,** Effect of the mutation on ZNF468 binding to the MER11 family. The violin plot on the right indicates the binding rate of ZNF468 to the MER11_2 subfamily with (left) and without (right) the 1258 C-to-T mutation. Statistical testing was conducted using the two-sided Mann–Whitney U test. ***p<0.001.

We focused on ERVs or long terminal repeats (LTRs) because HERVK, MER11, HERVH, and LTR7 were among the top results, in addition to LINE1. We observed many evasion candidates in MER11 family (**Fig. 3c**). MER11, the LTR of HERVK11, first emerged from the common ancestor of Catarrhines (29.4 MYA) and continued to transpose to the common ancestor of Homininae (9.1 MYA). Recent research has highlighted that the MER11 family comprises more subfamilies than previously identified^6^. Consequently, we reapplied our subfamily classification pipeline to the MER11 family and classified MER11 family into 6 subfamilies (**Supplementary Fig. 6a, b**). In the MER11 family, evasion from seven KRAB-ZFPs was observed, and the binding of these KRAB-ZFPs gradually disappeared over the course of MER11 evolution (**Fig. 3c** and **Supplementary Fig. 6c**). Among these KRAB-ZFPs, ZNF611, ZNF440, and ZNF808 emerged in the same era as their target MER11 subfamilies, whereas ZNF445, ZNF33A, ZNF468, and ZNF433 appeared in eras older than their target MER11 subfamilies. Furthermore, we identified the binding sites of ZNF468 and ZNF433, and found that the disappearance of binding was attributable to mutations or indels at their binding sites (**Fig. 3d** and **Supplementary Fig. 6d, e**). At the binding site of ZNF468, position 1258 was mutated during the evolution of MER11_2, preventing ZNF468 from binding to MER11 copies via a 1258 C-to-T mutation (**Fig. 3e**). These findings suggest that MER11 evolved to evade KRAB-ZFP suppression through point mutations and indels. Additionally, young MER11 subfamilies were targeted by ZNF525, ZNF578, and ZNF727, which complemented the loss of KRAB-ZFP binding (**Fig. 3c** and **Supplementary Fig. 6c)**. ZNF525 and ZNF727 bound to MER11 copies that were active in the same era as their emergence, suggesting that the MER11 family was targeted by the newly emerged KRAB-ZFPs after evasion from several KRAB-ZFPs. These findings illustrate the existence of an evolutionary arms race between the MER11 family and KRAB-ZFPs. A similar pattern was observed in the LTR5 family (**Supplementary Fig. 7**). In summary, these results suggest that evasion from KRAB-ZFPs occurred across a broader range of TE families than previously reported, leading to an evolutionary arms race with KRAB-ZFPs.

### Evolutionary arms races between ERVs and KRAB-ZFPs

ERVs exist within the genome either as proviruses or as solo LTRs^29^. ERVs are typically annotated by dividing them into LTRs and internal regions. Because ERVs expand through proviruses, to fully understand the evolutionary arms race between ERVs and KRAB-ZFPs, it is necessary to analyze ERVs not as separate LTRs and internal regions, but as proviruses. Therefore, we reconstructed proviruses from the TE annotation data (**Fig. 4a** and **Supplementary Fig. 8**). As a result of this reconstruction, we identified 5518 proviruses in 19 types of ERVs (**Fig. 4b** and **Supplementary Data 5**). We identified 48 LTRs and 82 internal regions as targets of KRAB-ZFP (**Supplementary Fig. 9a** and **Supplementary Data 2**). Interestingly, the binding rates of KRAB-ZFP varied not only in the LTR but also in the internal region between subfamilies (**Fig. 4c**). This trend was also observed in TRIM28 (**Supplementary Fig. 9b**). These findings suggest that alterations in sequences and KRAB-ZFP binding occur not only in LTRs but also in internal regions, implying that both regions had been involved in the evolutionary arms race with KRAB-ZFPs.

**Fig. 4.**
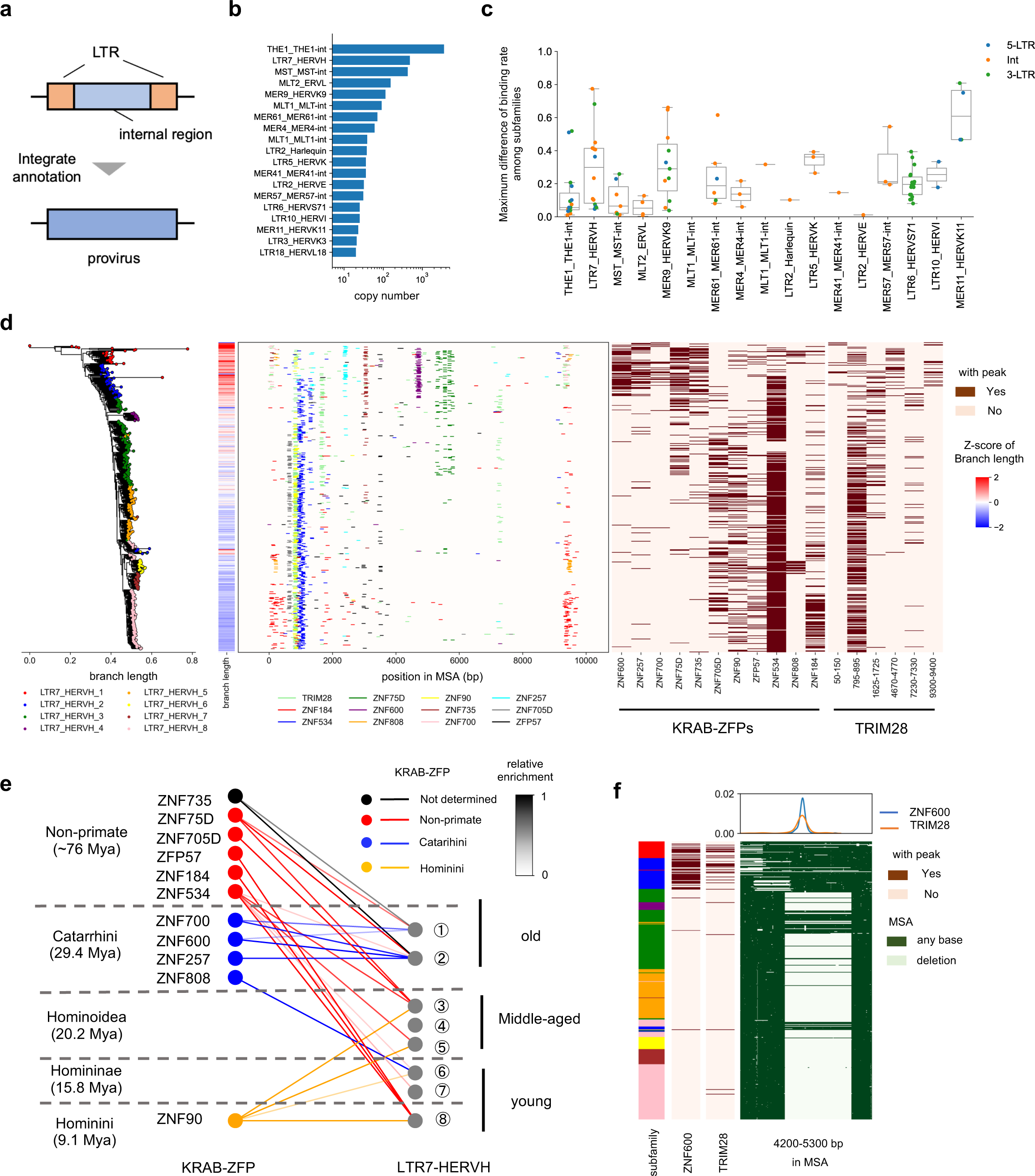
Evolutionary arms race with KRAB-ZFPs in full-length ERVs and LTR7-HERVH. **a,** Schematic representation of the reconstruction of proviruses by integrating separately annotated LTR and internal region sequences. **b,** The copy number of the provirus in each ERV family. **c,** Maximum difference in the binding rates of KRAB-ZFPs among subfamilies of each ERV family. Binding rates were estimated as the average proportion of ERV copies that overlapped with the KRAB-ZFP peaks. The maximum difference in the binding rate was defined as the difference between the maximum and minimum binding rates of the subfamilies in each KRAB-ZFP. The dotted colors indicate the region targeted by KRAB-ZFP, including the 5-LTR (blue), internal region (orange), and 3-LTR (green). **d,** Evolutionary arms race between LTR7-HERVH and the 11 KRAB-ZFPs. The phylogenetic tree and heat map of the branch length on the left indicate the phyletic relationship and insertion date, respectively. Peaks and binding patterns of KRAB-ZFPs and TRIM28 are shown at the center and right, respectively. **e,** Comparison of evolutionary ages between LTR7-HERVH subfamilies and KRAB-ZFPs. Dots indicate KRAB-ZFPs (left) and LTR7-HERVH subfamily (right). The lines between the dots show the relationship between the LTR7-HERVH subfamily and KRAB-ZFPs. The color and intensity of the lines indicate the evolutionary age of KRAB-ZFPs and the relative enrichment of their targets, respectively. **f,** Deletion of the ZNF600 binding site. The color bar (left) shows the LTR7-HERVH linkage in each copy, based on the phylogenetic tree. The two heat maps in the center indicate the binding of ZNF600 and TRIM28 between 4200 and 5000 bp, respectively. The top plot shows the positions of ZNF600 (blue) and TRIM28 (orange) peaks. The heatmap on the right shows the deletion of the sequence in each LTR7-HERVH copy. Flesh color indicates a deletion at each site.

Upon examining all proviruses in detail, remarkable characteristics of the evolutionary arms race with KRAB-ZFPs were observed in LTR7_HERVH (**Fig. 4d** and **Supplementary Fig. 10**) and THE1_THE1-int (**Supplementary Fig. 11**). We focused on LTR7_HERVH because LTR7_HERVH, an ERV essential for the pluripotency of hESCs^30,31^. LTR7_HERVH first emerged from the common ancestor of Catarrhines and remained active until the common ancestor of Homininae. Our pipeline classified LTR7_HERVH into 8 subfamilies, which is consistent with the subfamily classification reported in a previous study (**Supplementary Fig. 10a, b**)^32^. In LTR7_ HERVH, we observed 11 KRAB-ZFPs and 6 TRIM28 binding sites, most of which exhibited subfamily specificity (**Fig. 4d**). 5 KRAB-ZFPs targeted the old LTR7_HERVH subfamilies (LTR7_HERVH_1, 2) emerging from the common ancestor of Catarrhines, whereas the other KRAB-ZFPs targeted the middle-aged LTR7_HERVH subfamilies (LTR7_HERVH_3–5) and young LTR7_HERVH subfamilies (LTR7_HERVH_6–8) emerging from the common ancestor of Hominoidea and Homininae, respectively (**Fig. 4d** and **Supplementary Fig. 10c**). While the old LTR7-HERVH subfamilies and 3 of KRAB-ZFPs emerged in the same era (Catarrhini, 29.4 Mya), most KRAB-ZFPs targeting middle-aged and young LTR7_HERVH subfamilies existed before these subfamilies (**Fig. 4e**). Finally, young LTR7_HERVH was targeted by ZNF90, which emerged from the common ancestor of Homininae, suggesting a complementary response to the emergence of the new LTR7_HERVH subfamilies. Moreover, changes in the binding affinity of these KRAB-ZFPs were attributed to mutations and indels, similar to those observed in the MER11 family (**Fig. 4f** and **Supplementary Fig. 10d, e**). Specifically, the binding sites of ZNF600 were deleted in the LTR7_HERVH_3–8 subfamilies, along with the loss of its binding signal (**Fig. 4f**). This suggests that these middle-aged and young subfamilies evaded the regulatory control of ZNF600. These results suggest that LTR7_HERVH and some ERVs have undergone an evolutionary arms race with the KRAB-ZFP family.

### Evolutionary arms race shapes regulation landscape of LTR7_HERVH and nearby genes in hESCs

Finally, we aimed to reveal the relationship between the evolutionary arms race and co-option of LTR7_HERVH in the gene regulatory network of hESCs. Previous studies have shown that LTR7 is bound by pluripotency-related TFs (NANOG, KLF4, OCT4, SOX2, FOXP2, and FOXA1) and other TFs (FOXA2, GATA6, and EOMES)^2,32^. Therefore, we examined the binding patterns of KRAB-ZFPs, TRIM28, and these TFs, as well as the differences in chromatin state and expression between the LTR7_HERVH subfamilies in hESCs (**Supplementary Fig. 12a** and **Supplementary Table 1**)^23,33–35^. Consistent with the results of previous studies, young subfamilies, bound by KLF4, were highly expressed in hESCs (**Supplementary Fig. 12a**)^32,36^. In contrast, old and middle-aged subfamilies tended to overlap with the heterochromatin state (9_Het in ChromHMM 15 models) enriched with H3K9me3 modifications compared to the youngest LTR7_HERVH subfamily (**Supplementary Fig. 12a**). Although not statistically significant in some LTR7_HERVH subfamilies, LTR7-HERVH copies overlapping with 9_Het showed lower expression than those without an overlap with 9_Het (**Supplementary Fig. 12b**). To examine the roles of KRAB-ZFPs and TRIM28 in LTR7-HERVH expression in hESCs, we compared LTR7-HERVH expression in wild-type and TRIM28 knock out (KO) hESCs. Interestingly, the deficiency of TRIM28 induced the upregulation of older and middle-aged subfamilies and the downregulation of the youngest subfamily (**Fig. 5a**). The expression levels of these TFs did not change significantly (**Supplementary Fig. 12c**). Moreover, LTR7_HERVH copies overlapping with 9_Het were more highly upregulated than other LTR7_HERVH copies (**Fig. 5b**). Consistent with these findings, TRIM28 peaks were observed in LTR7_HERVH copies in hESCs (**Supplementary Fig. 12d**). In the old LTR7_HERVH subfamilies, several TRIM28 binding sites associated with KRAB-ZFPs, were observed. Together, these results suggest that the subfamily specific binding patterns of KRAB-ZFPs and TRIM28 formed through the evolutionary arms race also contribute to shaping the subfamily specific expression of LTR7_HERVH in hESCs.

**Fig. 5.**
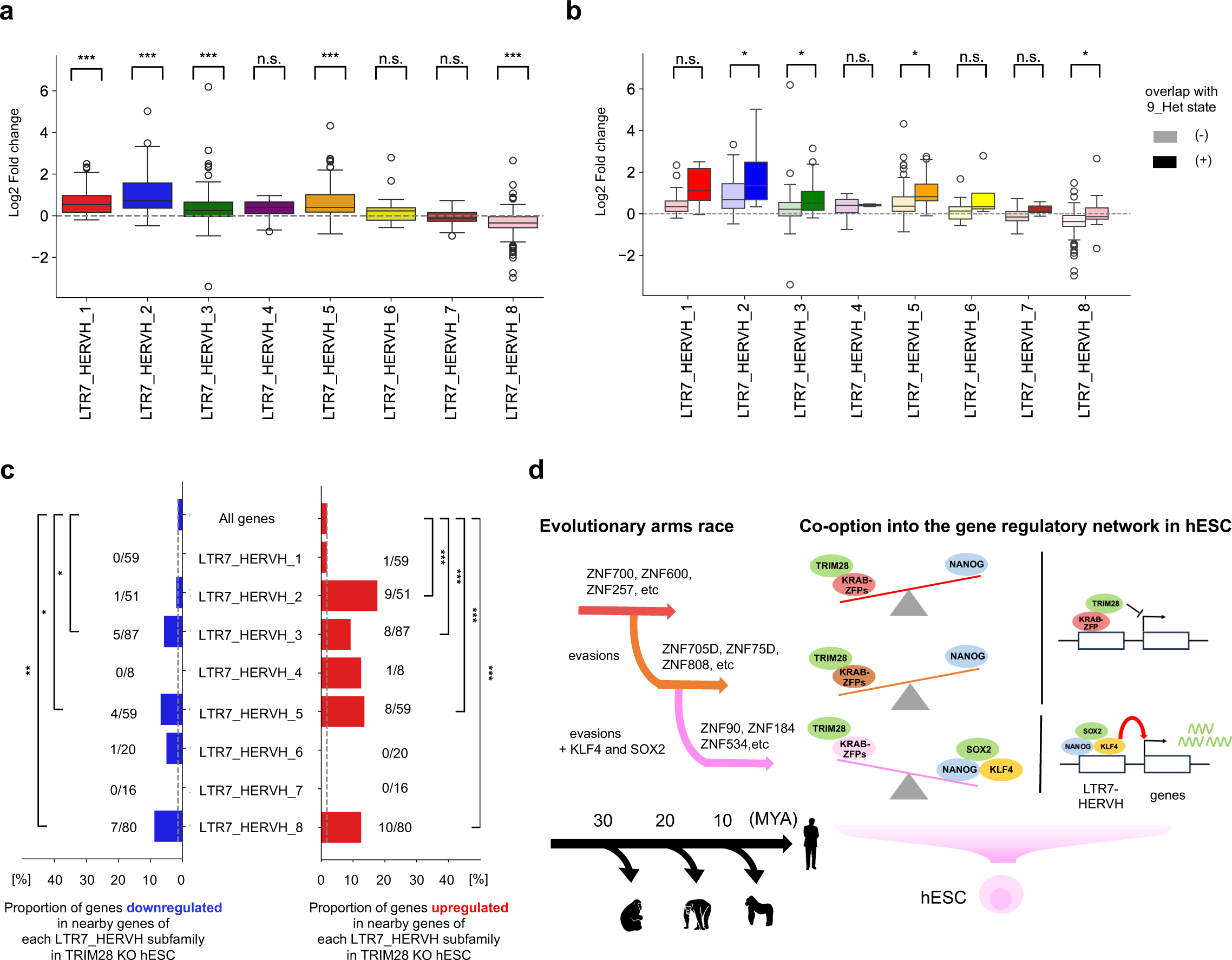
TRIM28 regulates the expression of LTR7-HERVH and nearby genes in hESCs. **a,** log2 fold change in the expression of LTR7-HERVH copies in TRIM28 knockout (KO) hESCs compared to wild-type (WT) hESCs for each LTR7-HERVH subfamily. Colors represent different subfamilies. Statistical analyses were performed using the two-sided Wilcoxon signed-rank test. P-values were adjusted for multiple testing using the Benjamini-Hochberg procedure ***FDR < 0.001; n.s., not significant. **b,** Relationship between perturbation due to TRIM28 deficiency and the heterochromatin state (9 Het) of LTR7-HERVH. The intensity of the color indicated an overlap with the heterochromatin state. Statistical testing was conducted using the two-sided Mann–Whitney U test. P-values were adjusted using the Benjamini-Hochberg procedure. *FDR< 0.05. **c,** Proportion of DEG in nearby genes of each LTR7_HERVH subfamily. Box plots on the left and right show the proportions of downregulated and upregulated genes among the nearby genes, respectively. Statistical analyses were conducted using the two-sided binomial test and compared with the proportion of DEGs in all genes. P-values were adjusted using the Benjamini-Hochberg procedure. *FDR<0.05, **FDR<0.01, ***FDR<0.001. **d,** Schematic model illustrating the effect of the evolutionary arms race between LTR7-HERVH and KRAB-ZFPs (left) on the regulatory landscape of LTR7-HERVH and nearby genes in hESCs (right). KRAB-ZFPs and TRIM28 silence older LTR7-HERVH subfamilies, whereas younger subfamilies are activated by pluripotent TFs. Silencing older subfamilies prevents the aberrant activation of nearby genes, whereas younger subfamilies function as cell type-specific regulatory elements.

Furthermore, many recent studies have demonstrated that derepression of TEs affects the expression of nearby genes^21,23,24^. Therefore, we verified the impact of alterations in LTR7_HERVH expression on the expression of nearby genes within 50 kb of the LTR7_HERVH copy. Consistent with our hypothesis, the expression of genes near the LTR7_HERVH copies was significantly altered more than that in all the genes (**Fig. 5c**). Even within 50 kb, differentially expressed genes (DEG) were closer to LTR7_HERVH copies than those that were not differentially expressed, consistent with the model in which derepressed TEs perturbed the expression of nearby genes (**Supplementary Fig. 12e**). Specifically, in the LTR7_HERVH_2 subfamily, which exhibited the greatest increase in expression as shown in **Fig. 5a**, the proportion of upregulated genes was significantly higher than that of all the genes (18% vs. 2%). However, the proportion of downregulated genes did not follow this pattern (2% vs. 2%), suggesting that derepression of LTR7_HERVH induced the upregulation of nearby genes. These findings suggest that KRAB-ZFPs/TRIM28 regulates gene expression via LTR7_HERVH. Our observations suggest that the binding pattern of KRAB-ZFPs/TRIM28 formed through the evolutionary arms race suppresses and modulates LTR7_HERVH subfamilies and nearby genes and contributes to the gene regulatory network in hESCs (**Fig. 5d**). In summary, our findings suggest that an evolutionary arms race with KRAB-ZFPs could lead to the co-option of LTR7-HERVH and KRAB-ZFPs in the gene regulatory network of hESCs.

## Discussion

In this study, we systematically investigated the evolutionary arms race relationship between TE families and KRAB-ZFPs (**Fig. 1**). We comprehensively identified the targets of KRAB-ZFPs and compared their evolutionary ages with those of their target TE subfamilies to characterize the chronological relationship between KRAB-ZFPs and TE subfamilies. (**Fig. 2**). Furthermore, we identified and characterized novel instances of evolutionary arms races in MER11, LTR5, LTR7_HERVH, and THE1_THE1-int (**Fig. 3, 4**). Remarkably, we also revealed that the evolutionary arms race in LTR7_HERVH was related to the regulation of LTR7_HERVH and nearby genes in hESCs (**Fig. 5**). In summary, our findings suggest that TEs, KRAB-ZFPs, and their host organisms have complex co-evolutionary relationships.

It has been hypothesized that KRAB-ZFPs have emerged to suppress newly emerging TE subfamilies^8,9,17^. Based on this hypothesis, we identified the TE targets of KRAB-ZFPs and obtained the evolutionary ages of both KRAB-ZFPs and TE subfamilies to re-examine the relationship between KRAB-ZFPs and TEs. Consistent with the results of previous studies, we observed that KRAB-ZFPs and their target TE subfamilies emerged concurrently (**Fig. 2b**). The expansion of KRAB-ZFPs and TE families occurred in the common ancestors of Eutherians, Simiiformes, and Catarrhines, re-emphasizing that KRAB-ZFPs engaged in evolutionary arms races through gene duplication during these periods (**Supplementary Fig. 4c**).

In addition to the gene duplication mechanisms discussed above, we observed that KRAB-ZFPs that emerged from the common ancestor with non-primates also targeted primate-specific TE subfamilies (**Fig. 2b** and **Supplementary Fig. 4f**). While this phenomenon has been reported in the youngest L1, our observations suggest that it is more universal across TE classes (**Supplementary Fig. 4e**)^8^. There are two possible explanations for this observation. First, ancient KRAB-ZFPs may have adapted to target and suppress newly emerged TE subfamilies. ZNF649, which emerged from the common ancestor of the Eutherians, has been reported to have acquired new ZF domains and sequence specificity during the evolution of primates, enabling ZNF649 to suppress young L1 elements, thus supporting this hypothesis^28^. The second scenario is the opposite of the typical evolutionary arms race in which younger primate-specific TEs have evolved to be bound KRAB-ZFPs. The uncontrolled expansion of TEs poses a threat to host survival and reproductive functions, suggesting a trade-off between TE proliferation and genetic persistence. Interactions with existing KRAB-ZFPs may have enabled the TEs to expand while maintaining host viability. However, these hypotheses require further investigations in future research.

Next, we identified many candidates for evasion events of TE families from KRAB-ZFP suppression (**Fig. 3a, b**). In addition to the young L1 family, which has previously been reported to evade KRAB-ZFP, many other events have been observed in ERVs. The loss of KRAB-ZFP binding in younger subfamilies was attributed to substitutions and indels in the TE sequences, which supports evasion from KRAB-ZFP over the evolution of TE families (**Fig. 3d, e** and **Fig. 4f**). As the deletion of KRAB-ZFP clusters in mice led to a slight increase in ERV insertions over the short span of a few generations, these events could have potentially induced the expansion of ERVs over millions of years^16^.

Additionally, we showed that several ERVs were targeted by the new KRAB-ZFPs following their evasion from KRAB-ZFPs. Specifically, we demonstrated that, in the MER11 family, after the binding of 7 KRAB-ZFPs targeting old MER11 copies disappeared, three newly emerged KRAB-ZFPs alternatively targeted younger copies (**Fig. 3c** and **Supplementary Fig. 6c**). This retargeting by KRAB-ZFPs represents a complementary response to the emergence and expansion of young TE subfamilies that have evaded KRAB-ZFPs, suggesting the occurrence of evolutionary arms races between these ERVs and KRAB-ZFPs. It is important to note that, in many analyses, we limited ourselves to TE families that were active across multiple eras to examine the chronological relationship between the emergence of TE families and KRAB-ZFPs. In TE families that expanded over a short period, we were unable to examine TE events or evolutionary arms races. Nevertheless, our findings revealed that evolutionary arm races with KRAB-ZFPs occurred in more TE families than previously reported.

Furthermore, we demonstrated that the expression differences of the LTR7_HERVH subfamily in hESCs were determined not only by transcription factors but also by KRAB-ZFPs/TRIM28. It has been suggested that the young subfamilies of LTR7_HERVH are specifically activated in hESCs because the transcription factors KLF4 and SOX2 bind specifically to these subfamilies^32^. However, our analysis showed that even older subfamilies could be activated in hESCs when the suppression by TRIM28 was inhibited (**Fig. 5a, b**). Additionally, our data suggest that young LTR7_HERVH subfamilies can be activated in hESCs despite being targeted by TRIM28 to the same extent as older subfamilies (**Supplementary Fig. 12d**). This phenomenon is likely due to KLF4 and SOX2, which bind to these young subfamilies and activate transcription, even within the heterochromatin, functioning as pioneer transcription factors^32,37^. In summary, our data suggested that the subfamily specific expression patterns of LTR7_HERVH were formed by a balance between activation by transcription factors and suppression by KRAB-ZFPs/TRIM28 (**Fig. 5d**).

Finally, we investigated the effect of TRIM28 deficiency on the expression of genes near the derepressed LTR7_HERVH. Importantly, we discovered that derepression of older LTR7_HERVH subfamilies tended to promote the upregulation of nearby genes (**Fig. 5c**). This finding suggests that in the absence of TRIM28 suppression, the old LTR7_HERVH subfamilies can also act as cis-regulatory elements that influence nearby genes. Young LTR7__ HERVH subfamilies, which are specifically activated in hESCs, play a crucial role in maintaining pluripotency in hESCs, likely through the regulation of neighboring gene expression. Overall, our data suggest that KRAB-ZFPs/TRIM28 modulates not only the expression patterns of LTR7_HERVH, but also those of genes near the LTR7_HERVH copies (**Fig. 5d**).

This study has three limitations. The first was the inability to infer causality during the evasion of KRAB-ZFPs. Although our findings and those of previous studies support the concept of TE evasion, we have not examined whether the loss of KRAB-ZFP binding in younger subfamilies is due to random drift or selective pressure. In a future study, we need to examine whether evasion from KRAB-ZFPs accelerates the proliferation of TEs within the host genome. The second limitation is the possibility that the binding patterns of human KRAB-ZFPs do not necessarily reflect those of their ancestors. Given the rapid evolution of KRAB-ZFP genes, it is possible that some ancestral KRAB-ZFPs were lost or optimized for co-option in the gene regulatory network. Additionally, old TE families have lost KRAB-ZFP-binding sites because of the accumulation of mutations. This led to a decrease in KRAB-ZFP binding affinity, making it difficult to detect in analyses. Although the binding patterns of KRAB-ZFPs and TRIM28 in humans may retain aspects of their ancestral state, they may represent optimized or attenuated conditions. Third, the present study did not consider suppression mechanisms other than KRAB-ZFPs and TRIM28. Mechanisms that suppress TE transposition in host organisms are robust and complementary. For example, it has been reported that the youngest L1 is not repressed by TRIM28 but is instead repressed by DNA methylation associated with PIWI-interacting RNA^38^. Therefore, these mechanisms may complement the gaps in the evolutionary arms race with KRAB-ZFPs. Cross-species analyses that include various transposition suppression mechanisms are necessary to fully elucidate the evolutionary arms race with KRAB-ZFPs.

Despite these limitations, we have demonstrated that the evolutionary arms race between TEs and KRAB-ZFPs is not limited to LINE1 but is universally observed across a wide range of TEs and that such evolutionary arms races may have influenced the evolution of the host’s gene expression regulatory network. This study provides new insights into the complex coevolutionary relationships among TE families, KRAB-ZFPs, and host gene regulatory networks.

## Methods

### Processing of ChIP-seq and ChIP-exo data

Raw reads from ChIP-seq and ChIP-exo were obtained from the NCBI GEO and Encyclopaedia of DNA Elements (ENCODE) databases (**Supplemental Table 1**)^39,40^. The ChIP-seq dataset for KRAB-ZFPs included 1051 samples, encompassing 361 KRAB-ZFP genes, from five previous large-scale studies on KRAB-ZFPs (GSE58341, GSE76496, GSE78099, GSE120539, GSE200964)^8,9,17,41,42^ and the ENCODE. The TRIM28 dataset consisted of 35 samples from 14 studies and the ENCODE. Datasets for TFs (NANOG, KLF4, SOX2, POU5F1, EOMES, FOXA1, FOXA2, and GATA6) in hESCs and differentiated cells were derived from GSE61475^33^ and GSE130417^23^. Raw reads were trimmed using fastp (ver.0.2.3) and mapped to GRCh38/hg38 using bowtie2 (ver.2.4.4) in an end-to-end –sensitive mode^43,44^. Unmapped reads were removed, and SAM files were converted to BAM using SAMtools (ver1.9)^45^. Multiple mapped reads were retained for analysis. PCR duplicates were removed only from ChIP-seq samples using Picard (ver.2.9.2) and SAMtools, because keeping them in ChIP-exo samples was recommended^8,46^. Peak calling was performed using MACS2 (ver.2.2.7.1) with the options –keep-dup all -q 0.05^47^. The peaks were filtered using the following criteria: (1) does not overlap with the blacklist regions defined by the ENCODE (ENCFF419RSJ), (2) signal value > 2, (3) score > 50, and (4) peak length < 1000 bp. The ChIP-seq samples without inputs were also processed using this pipeline.

### Subfamily classification and genome annotation of TE families and proviruses

Genomic annotation data for the TE families were obtained using a two-step process (**Supplementary Fig. 3**). In the first step, subfamily classification was performed using the genomic annotation data of TEs. The GRCh38/hg38 genome was annotated using RepeatMasker with consensus sequences of the TE subfamilies downloaded from Dfam^48^. Fragmented repeats were reconstructed using OneCodeToFindThemAll.pl^49^. All extracted repeats were aligned to these consensus sequences using MAFFT (ver.7.520) with the options --addfragments --keeplength --retree 2 --mapout^50^. To filter out fragmented and truncated repeats, those that did not meet the following criteria were excluded from the analysis: alignment to the consensus sequence was less than 80% (or 60% if the consensus sequence length exceeded 4000 bp); (2) more than 20% of the repeat were not aligned to the consensus sequence; and (3) the substitution rate exceeded 0.4 times per base pair. TE that met these criteria were defined as full-length copies. TE families with more than 20 full-length sequences at the family level were aligned using MAFFT with the option --auto. To reduce the computational load on TE families with more than 10,000 full-length copies, 10,000 full-length copies were randomly sampled for analysis. Positions with more than 99% of the gaps in the mulple sequence alignment (MSA) were removed. Phylogenetic trees were constructed using iqtree2 with the options -m MFP -bb 1000 -alrt 1000^51–53^. The maximum likelihood distance matrix presented in the mldist file was dimensionally reduced using UMAP^54^. The latent space obtained from UMAP was clustered using K-means (k=2–20). The optimal number of clusters was manually determined based on the number of clusters with the highest average silhouette coefficients. Of the 173 TE families without subfamily classification and with more than 20 full-length copies, new clustering was adopted for 59 families with an average silhouette coefficient exceeding 0.6; these clusters were defined as new subfamilies. From the alignment data, the consensus sequences for each subfamily were reconstructed based on the majority-rule consensus. If more than 50% of the bases were present, a specific base was used; otherwise, ’N’ was assigned. The consensus sequences of the original family were replaced with those of the newly defined subfamilies.

Second, the GRCh38/hg38 genome was re-annotated using new consensus sequences. The process was performed using the same pipeline as in the first step. The full-length sequences were defined according to the first two criteria established in the first step. To eliminate the misannotation of Alu sequences, the third criterion was specifically applied to the SVA family. In all steps, repeats located on the Y chromosome, contigs, and unmapped regions (described later) were removed.

The genome annotation and subfamily classification of the MER11 family were constructed based on the annotation of 2^nd^ step due to the heterogenicity of the MER11 subfamily highlighted by previous research and the redundancy of the annotation between the MER11A and MER11C subfamilies^6^. Duplications in the annotation of MER11 copies longer than 700bp were examined using Bedtools intersect. Annotations of the MER11 copies that were longer and had higher scores were retained. A total of 2239 MER11 copies were used in the analyses. Phylogenetic tree construction and subfamily classifications were performed using the pipeline described above.

The LTRs and internal regions constituting proviruses were determined based on a previous study and Dfam^26,55^. The LTRs and internal regions were assembled using OneCodeToFindThemAll.pl. The overlaps between the assembled annotations and the annotations obtained in the first step were examined using Bedtools intersect. Assembled sequences meeting the following criteria were defined as full-length: (1) overlap of one full-length internal region and two full-length LTRs; (2) both LTRs belonging to the same subfamily; and (3) the internal region flanked by the two LTRs. Provirus families were classified into subfamilies using the process outlined in the above subfamily classification pipeline.

### Definition of KRAB-ZFP targets

The target of each KRAB-ZFP was determined using a binomial test, as described in a previous study^9^. Overlaps between the summits of the KRAB-ZFP peaks and TE subfamilies were quantified using Bedtools intersect. The expected overlap probability was estimated by dividing the total length of each TE subfamily by the effective genomic length, which was defined as the total genome length, excluding the Y chromosome, contigs, and unmapped regions. Unmapped regions were inferred from 405 ChIP-seq samples. The coverage for each 100 bp bin was quantified using Bedtools coverage. Regions with zero coverage across all the ChIP-seq samples were defined as unmapped regions. The q value was calculated for each KRAB-ZFP using the Benjamini-Hochberg procedure. Targets were considered significant if the q value was < 0.05. A target was defined as the primary target if the quotient of the -log10 q value divided by the maximum value of the -log10 q value exceeded 0.9; otherwise, it was defined as a secondary target. In addition to the primary targets, secondary targets were included in the analysis if they met the following criteria: (1) -log10 q value > 10; (2) ratio > 2; (3) more than 5 peaks overlapping with full-length targets; and (4) peaks observed in more than 10% of the full-length targets. When a KRAB-ZFP targeted at least one subfamily belonging to a certain family, it was defined as a KRAB-ZFP targeting that family.

The full-length ERV targets of each KRAB-ZFP were determined for the 5-LTR, internal region, and 3-LTR. Statistical testing was performed using the same pipeline as the TEs, and KRAB-ZFP-ERV subfamily associations that met the following criteria were used for analyses: (1) -log10 q value > 10; (2) ratio > 2; (3) more than 5 peaks overlapping with full-length targets; and (4) peaks observed in more than 10% of the full-length targets.

### Evolutionary age inference of TE subfamilies, proviruses, and KRAB-ZFPs

The evolutionary ages of the TE subfamilies and TE copies were estimated by liftover to the genomes of the 39 species (**Supplementary Table 2**). The chain files for the liftover were downloaded from UCSC^56^. All TE copies were lifted to other species using LiftOver -minMatch=0.5. The evolutionary age of the TE copies was defined as the oldest era in which they could be lifted to at least three species, or more than half of the species within the same branch. The evolutionary age of TE subfamilies was defined as the oldest era in which more than 10% of the full-length copies first appeared. Because proviruses can lose their internal region and one LTR due to homologous recombination, the evolutionary ages of proviruses were defined using the evolutionary ages of the 5-LTR rather than that of the full-length provirus.

The evolutionary ages of the KRAB-ZFPs were obtained from two previous studies^8,9^. The evolutionary ages of KRAB-ZFPs were primarily determined based on the estimated ages of de Tribolet-Hardy et al^9^. Missing values were supplemented using data from Imbeault et al^8^.

### Screening of TE evasion from KRAB-ZFPs

For each significant combination of TE families and KRAB-ZFPs, differences in the binding rate of KRAB-ZFPs among all pairs of TE subfamilies were examined. The binding rate was calculated as the mean proportion of overlap in the ChIP-seq samples. Statistical testing was performed using Tukey’s Honest Significant Difference test. A candidate for an evasion event of the TE family from KRAB-ZFP was defined as having at least one pair of subfamilies that met the following criteria: (1) adjusted p-value < 0.05, (2) difference in binding rates > 0.1, (3) decrease in binding rate in the younger subfamily, and (4) KRAB-ZFP being older than at least one TE subfamily (**Supplementary Data 5**).

### Phylogenetic analyses of TE families

An unrooted phylogenetic tree was constructed using iqtree2 with the same options employed for subfamily classification. The oldest subfamily was defined as the subfamily with the oldest evolutionary age and the longest branch length. As the root of the phylogenetic tree, the copy that emerged during the evolutionary age of the oldest subfamily and had the longest branch length was selected.

### MSA analyses of TE families

Consensus sequences of the TE subfamilies were aligned using MAFFT with the option --auto. All copies were aligned to the MSA of consensus sequences using MAFFT with the options --addfragments --keeplength --retree 2 --mapout. The positions of the ChIP-seq peaks in hg38 were converted to positions in MSA using map files.

### De novo motif analyses and identification of KRAB-ZFP binding sites

For the de novo motif analysis, all peaks overlapping with TEs, satellites, and simple repeats were excluded. The top 500 peaks with the highest signal values were used. If there were fewer than 500 peaks, all the peaks that did not overlap with the repeats were used. Sequences 250 bp upstream and downstream of the peak summits were used as inputs. De novo motifs for each KRAB-ZFP experiment were generated using MEME (ver.5.3.0) with the options meme-chip -dna -minw 6 -maxw 30 -meme-nmotif 5 -meme-p 8 -meme-mod zoops^57^. The positions of the motifs within the peaks were identified using FIMO with options – thresh 0.0001 –no-qvalue^58^. Motif positions were aligned on the MSA. The discovery rate within the peaks was calculated for each motif position in MSA. Positions with a discovery rate of 50% or higher were defined as binding sites.

### Processing of RNA-seq data and DEG analyses

Raw reads were downloaded from the GSE99215 dataset (**Supplementary Table 3**)^15^. The raw reads were trimmed using fastp and mapped to hg38 using STAR (ver. 2.7.8) with the options –outSAMtype BAM SortedByCoordinate – outFilterMultimapNmax 10000000 –outSAMmultNmax 1 –outMultimapperOrder Random^59^. Read counts of genes and LTR7-HERVH were separately counted using featureCounts (ver.2.0.1)^60^. Gene annotation of hg38 (GRCh38.13, release 40) was downloaded from GENCODE^61^. Differentially expressed gene (DEG) analyses between wild-type and TRIM28 KO hESCs were performed using DESeq2^62^. The read counts of LTR7-HERVH were normalized using the size factor estimated using with DESeq2.

Nearby genes were defined as genes whose transcription start sites (TSS) were within 50 kb of the LTR7_HERVH copies because previous studies have reported that DEGs were enriched within 50 kb of perturbed TE copies^23^. The TSS of the genes were extracted from the gene annotation. The distance between the gene and the LTR7_HERVH copy was defined as the shortest distance from either end of the LTR7_HERVH copy to the TSS. If the TSS overlapped with the LTR7_HERVH copies, the distance was defined as zero. To avoid the effects of pseudogenes, genes annotated as “protein_coding,” “lncRNA,” and “miRNA” were used for analyses.

### Chromatin state analyses with ChromHMM

The chromatin state of the hESCs (E003) was downloaded from Roadmap Epigenetics^34,35^. A Core15-state model lifted to hg38 was used. The overlap between chromatin states and LTR7-HERVH copies was obtained using bedtools intersect.

### Statistical analyses

Statistical analyses were performed using SciPy (ver.1.12.0) and statsmodels (ver.0.13.2)^63,64^. Multiple testing was performed using the Benjamini-Hochberg procedure.

### Code availability

The computational codes and processed data for reproduction are attached as a zip file. The code will be disclosed on Github (https://github.com/hmdlab/Evolutionary-Arms-Race) before publication.

## Acknowledgements

We would like to thank all co-authors, Dr. Jumpei Ito and Prof. Michiaki Hamada. We thank the members of Trono lab, Laboratory of Virology and Genetics at EPFL, for their discussions and insights that enriched our research. We also thank Junna Kawasaki for meaningful comments regarding this study. We thank all contributors to the public databases and data used in this study. We also extend our gratitude to all contributors who developed and maintained the bioinformatics tools and packages utilized in the present study. Computations were partially performed on the NIG supercomputer at ROIS National Institute of Genetics.

This study was supported in part by JST SPRING (JPMJSP2128, to Masato Kosuge).

## Author Contributions

Masato Kosuge performed all bioinformatics analyses.

Masato Kosuge, Jumpei Ito, and Michiaki Hamada designed the analyses and interpreted the results.

Masato Kosuge, Jumpei Ito and Michiaki Hamada wrote the original manuscript. All authors reviewed the manuscript.

## Conflict of interest

All authors declare that they have no conflict of interest.

## Figure legends

**Supplementary Table 1.** Information on ChIP-seq of KRAB-ZFPs, TRIM28, and TFs.

**Supplementary Table 2.** Species used to infer the evolutionary ages of TE copies and subfamilies.

**Supplementary Table 3.** Information on RNA-seq data

**Supplementary Data 1.** Metadata and evolutionary age of TE families and subfamilies.

**Supplementary Data 2.** KRAB-ZFP primary and secondary targets of the TE and full-length ERV subfamilies.

**Supplementary Data 3.** Metadata and evolutionary age of KRAB-ZFPs.

**Supplementary Data 4.** Candidates of TE evasion from KRAB-ZFPs.

**Supplementary Data 5.** Metadata and evolutionary age of full-length ERV families and subfamilies.

**Supplementary Fig. 1. Dataset of KRAB-ZFP ChIP-seq and ChIP-exo.**

**a,** Proportion of KRAB-ZFPs with ChIP-seq in public databases.

**b,** Dataset of KRAB-ZFP ChIP-seq. Upset plot displays the number of KRAB-ZFPs included in each ChIP-seq sample and the intersections among these datasets.

**Supplementary Fig. 2. Evaluation of the novel subfamily classification pipeline.**

**a,** Proportion of TE family without subfamily classification.

**b,** Clustering similarity (y-axis) of 162 TE families between annotations using RepeatMasker with TE consensus sequences in Dfam and our pipeline. Clustering similarity was defined using the ARI. The x-axis represents the average silhouette coefficient for each TE family. The colors of each dot indicate the TE classes, including ERVs (blue), LINEs (orange), SINEs (green), retroposons (red), and DNA transposons (purple).

**c,** Average silhouette coefficient (y-axis) of each TE family without subfamily classification.

**d,** Subfamily classification of LTR36 using our pipeline and consistency with the phylogenetic tree of LTR36. The UMAP plot on the left represents the latent space derived from the maximum likelihood distance matrix of LTR36. The color of each dot in the plot and each external branch in the phylogenetic tree indicates the subfamily of each copy.

**Supplementary Fig. 3. Detail of the subfamily classification pipeline and genome annotation.**

Schematic of the pipeline for subfamily classification of TE families (1st step) and genome annotation (2nd step). In the 1^st^ step, we annotated the hg38 genome using consensus sequences obtained from Dfam. Subsequently, repeats were extracted from the hg38 genome, filtered, and aligned. Using the alignment data, we constructed phylogenetic trees and calculated the maximum likelihood distance matrix using the iqtree2. We then performed dimension reduction on the distance matrix using UMAP and k-means clustering. Finally, we constructed new subfamily classifications and consensus sequences for 59 TE families that met these criteria. Original consensus sequences were replaced with consensus sequences. In the 2^nd^ step, a new set of consensus sequences was used to annotate the hg38 genome, which was then used for subsequent analyses. Please refer to the Methods section for further details.

**Supplementary Fig. 4. Chronological relationship between TE families and KRAB-ZFPs.**

**a,** Proportion of KRAB-ZFP targeting TEs.

**b,** Proportion of the TE subfamily (left) and TE family (right) targeted by KRAB-ZFPs.

**c,** Evolutionary ages of KRAB-ZFPs (blue) and TE subfamilies(orange). The x-axis indicates the evolutionary age (MYA) and the y-axis indicates the probability density estimated by kernel density estimation.

**d,** Comparison of evolutionary ages between the TE subfamily (x-axis) and KRAB-ZFPs (y-axis) in 1804 primary and secondary KRAB-ZFP-TE subfamily associations.

**e,** Comparison of evolutionary ages of ERVs (left) and LINEs (right) with KRAB-ZFPs. Contour lines were estimated from 268 and 208 primary KRAB-ZFP-TE subfamily associations in ERVs and LINEs, respectively.

**f,** Preference for non-primate (left) and primate (right) KRAB-ZFPs for the evolutionary ages of the target TE subfamilies. Statistical tests were conducted using the chi-square test. ***p<0.001.

**Supplementary Fig. 5. Evasions of L1P family from KRAB-ZFPs**

**a,** Evasion of L1P_5end from KRAB-ZFPs. The phylogenetic tree indicated a phyletic relationship between the L1P_5end copies. Heatmap plots of branch length and liftover indicate the insertion date of each L1P_5end copy. The heatmap plot on the right shows the binding profile of KRAB-ZFPs that may have been evaded by L1P_5end family.

**b,** Evasion of the L1P family from ZNF765, ZNF649, and ZNF93 through point mutations and deletions. Color bars indicate the L1P_5ned subfamilies. The heatmap on the left indicates the overlap of KRAB-ZFP peaks with each L1P_5end copy. The sequence logs of ZNF765 and ZNF649 represent the de novo motif of KRAB-ZFP (upper panel) and the consensus sequence derived from L1P_5end copies with KRAB-ZFP peaks (lower panel), respectively. The ZNF93 plot shows the distribution of ZNF93 peaks. The heatmap on the right represents a match with the consensus sequence or deletion in each position and copy.

**Supplementary Fig. 6. Subfamily classification of MER11 and the evasion from KRAB-ZFPs via point mutations and deletions**

**a,** Subfamily classification of the MER11 family. The plot shows the latent space of the MER11 family obtained through dimensionality reduction using UMAP. Dots and colors represent copies and subfamilies, respectively.

**b,** Relationship between known subfamilies and subfamily classification using our pipeline. The color of each dot on the left represents a known subfamily. The heatmap and color bar represent the proportion of known subfamilies within each subfamily as determined by our pipeline. The heatmap represents the overlap between known subfamilies and each subfamily as determined by our pipeline. **c,** Binding profile of KRAB-ZFPs to MER11 family. Color bar on the left represents the MER11 linkage in each copy. The heatmap shows the overlap of KRAB-ZFP peaks with each MER11 copy.

**d,** Evasion from ZNF433 via mutations. Sequence logos indicate the de novo motif of ZNF433 (top) and the sequence of ZNF468 binding site in the MER11 subfamily. Letters indicate the proportion of bases in each position.

**e,** Evasion from ZNF468 and ZNF433. Color bars indicate the MER11 subfamilies. The heat map on the left indicates the overlap of KRAB-ZFP peaks with each copy. The sequence logs at the top represent the de novo motif of KRAB-ZFP and the consensus sequence derived from MER11 copies with KRAB-ZFP peaks. The heatmap on the right shows a match between the consensus sequence in each position and the copy.

**Supplementary Fig. 7. Evolutionary arms race between LTR5 and KRAB-ZFPs.**

The phylogenetic tree indicates a phyletic relationship between the LTR5 copies. Heatmap plots of branch lengths and liftovers indicate the insertion date of each LTR5 copy. The heatmap plot on the right shows the binding profile of KRAB-ZFPs that may have evaded or targeted LTR5 subfamilies with -log10 FDR > 50.

**Supplementary Fig. 8. Detail of the pipeline for whole genome provirus annotation**

Schematic of the pipeline for the identification of proviruses and subfamily classifications. The LTRs and internal regions were assembled using OneCodeToFindThemAll.pl. Proviruses were extracted by comparing with the TE annotations obtained in the 1^st^ step (See Supplementary Fig.3 and Methods). For each ERV family, an MSA and phylogenetic tree of proviruses were constructed. Subsequently, subfamily classification was performed using UMAP and K-means.

**Supplementary Fig. 9. Difference of KRAB-ZFPs/TRIM28 binding in LTRs and Internal regions.**

**a,** Number of KRAB-ZFP targets in LTRs and internal regions of full-length ERVs. **b,** Maximum difference in TRIM28 binding rates among the subfamilies. The colors of the dots indicate the regions, including the 5-LTR (blue), internal region (orange), and 3-LTR (green).

**Supplementary Fig. 10. Subfamily classification of LTR7_HERVH family and the evasion from KRAB-ZFPs.**

**a,** Subfamily classification of the LTR7_HERVH family. The plot shows the latent space of the LTR7_HERVH family obtained through dimensionality reduction using UMAP. Dots and their colors represent the copies and their respective subfamilies.

**b,** Comparison of our dataset with that of Carter et al. The left Venn diagram shows the overlap between the two datasets. Heatmap and color bar show the proportion of LTR7 subfamilies identified by Carter et al. within each subfamily, as determined by our pipeline.

**c,** Insertion date of the LTR7_HERVH subfamily. The heat map on the left shows the proportion of LTR7_HERVH copies that emerged in each era. The box plot on the right shows the branch lengths of each LTR7_HERVH subfamily.

**d,** Evasion from ZNF75D. The color bar indicates the LTR7_HERVH subfamily. The heatmap on the left indicates the overlap of ZNF75D peaks with each copy. The sequence logs at the top represent the de novo motif of ZNF75D and the consensus sequence derived from LTR7_HERVH copies with ZNF75D peaks. The heatmap on the right shows a match between the consensus sequence in each position and the copy.

**e,** Effect of the mutation on ZNF75D binding to the LTR7_HERVH family. The violin plot on the right indicates the binding rate of ZNF75D to the LTR7_HERVH_1–4 subfamily with and without the 5557 mutations. Statistical testing was conducted using the two-sided Mann–Whitney U test. ***p<0.001.

**Supplementary Fig. 11. Evolutionary arms race between THE1_THE1-int and KRAB-ZFPs.**

**a,** Evolutionary arms race between THE1_THE1-int and KRAB-ZFPs. The phylogenetic tree indicates a phyletic relationship between the THE1_THE1-int copies. Heatmap plots of the branch length and the plot on the lower left indicate the insertion date of each copy. The plot on the right shows ChIP-seq peaks of TRIM28 and KRAB-ZFPs. PRDM9 was removed because it did not have a consistent binding site.

**b,** Comparison of evolutionary ages between THE1_THE1-int subfamilies. The heat map shows the proportion of THE1_THE1-int copies that emerged in each era.

**c,** Binding profile of KRAB-ZFPs in the THE1_THE1-int family. The color bar on the left represents the THE1_THE1-int linkage in each copy. The heatmap shows the overlap of KRAB-ZFP peaks with each THE1_THE1-int copy.

**Supplementary Fig. 12 Effect of TRIM28 deletion in hESCs on the expression of LTR7_HERVH, nearby genes, and TFs.**

**a,** Comparison of the binding patterns of TFs and chromatin state and expression in hESCs between the LTR7-HERVH subfamilies. Heatmaps show the proportion of LTR7_HERVH copies with overlapping TF peaks or chromatin states. The box plot on the right indicates the normalized read counts of LTR7-HERVH copies. **b,** Relationship between LTR7-HERVH expression and heterochromatin state. The y-axis indicates the normalized read count of LTR7-HERVH copies. Darker and lighter colors indicate whether there was an overlap with the heterochromatin state (9_Het) in hESCs (E003). Statistical testing was conducted using the two-sided Mann-Whitney U test. P-values were adjusted using the Benjamini-Hochberg procedure. *FDR<0.05.

**c,** Effect of TRIM28 KO on TF expression. The white bars and black bars represent the means ± SD of the normalized read counts of TFs in TRIM28 wild and KO hESCs, respectively. Statistical testing was performed using DESeq2. n.s. not significant.

**d,** Histogram of TRIM28 peak summits in each LTR7_HERVH group of hESCs. The number of copies in each LTR7_HERVH group is indicated by “N=”. Arrows and gene symbols of KRAB-ZFPs above the histogram indicate TRIM28 peaks related to KRAB-ZFPs.

**e,** Distance distribution of DEGs and non-DEGs within 50 kbp of LTR7_HERVH copies. The x- and y-axes represent the distance classes from the LTR7_HERVH copies and the proportion of distance classes in nearby genes, respectively. “wild” and “TRIM28” represent the genes upregulated in TRIM28 wild-type and KO hESCs, respectively. Statistical analysis was conducted using the two-sided binomial test to compare the proportions of non-DEGs. P-values were adjusted using the Benjamini-Hochberg procedure. **FDR<0.01.

## Notes

### Competing Interest Statement

The authors have declared no competing interest.

