## Supplementary Figure 1-12 for "Landscape of Evolutionary Arms Races between Transposable Elements and KRAB-ZFP Family"

a

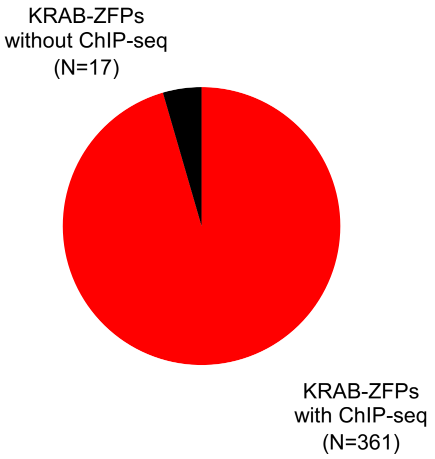


b

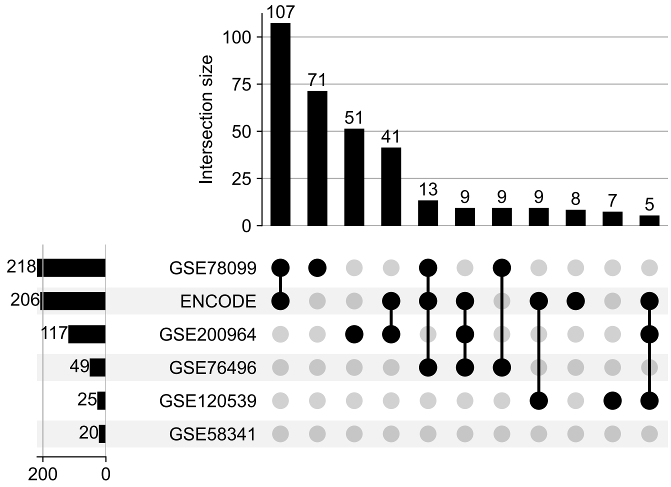


**Supplementary Fig. 1. Dataset of KRAB-ZFP ChIP-seq and ChIP-exo.**
**a,** Proportion of KRAB-ZFPs with ChIP-seq in public databases.
**b,** Dataset of KRAB-ZFP ChIP-seq. Upset plot displays the number of KRAB-ZFPs included in each ChIP-seq sample and the intersections among these datasets.

Supplementary Figure 2

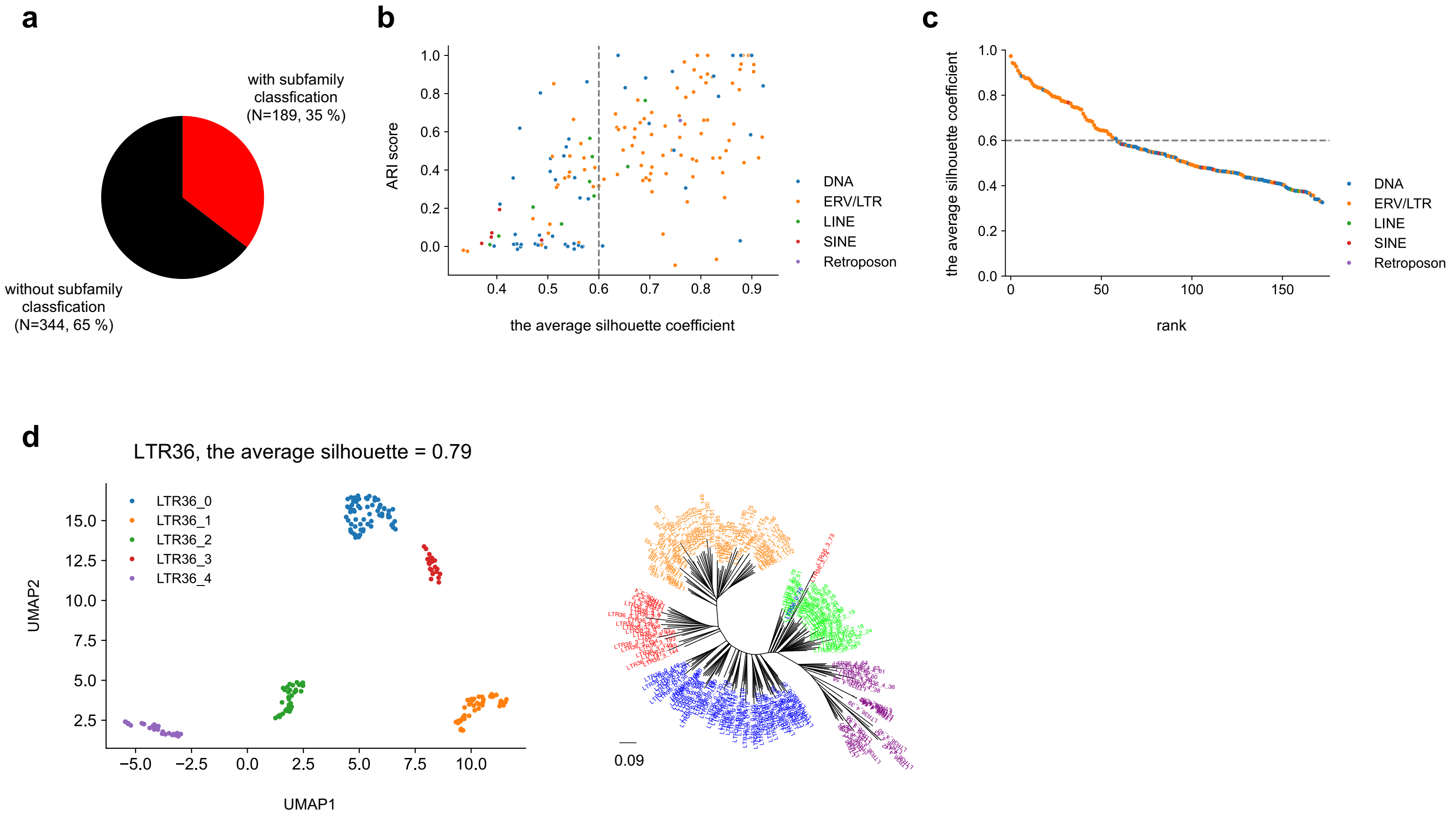


**Supplementary Fig. 2. Evaluation of the novel subfamily classification pipeline.**
**a,** Proportion of TE family without subfamily classification.
**b,** Clustering similarity (y-axis) of 162 TE families between annotations using RepeatMasker with TE consensus sequences in Dfam and our pipeline. Clustering similarity was defined using the ARI. The x-axis represents the average silhouette coefficient for each TE family. The colors of each dot indicate the TE classes, including ERVs (blue), LINEs (orange), SINEs (green), retroposons (red), and DNA transposons (purple).
**c,** Average silhouette coefficient (y-axis) of each TE family without subfamily classification.
**d,** Subfamily classification of LTR36 using our pipeline and consistency with the phylogenetic tree of LTR36. The UMAP plot on the left represents the latent space derived from the maximum likelihood distance matrix of LTR36. The color of each dot in the plot and each external branch in the phylogenetic tree indicates the subfamily of each copy.

Supplementary Figure 3

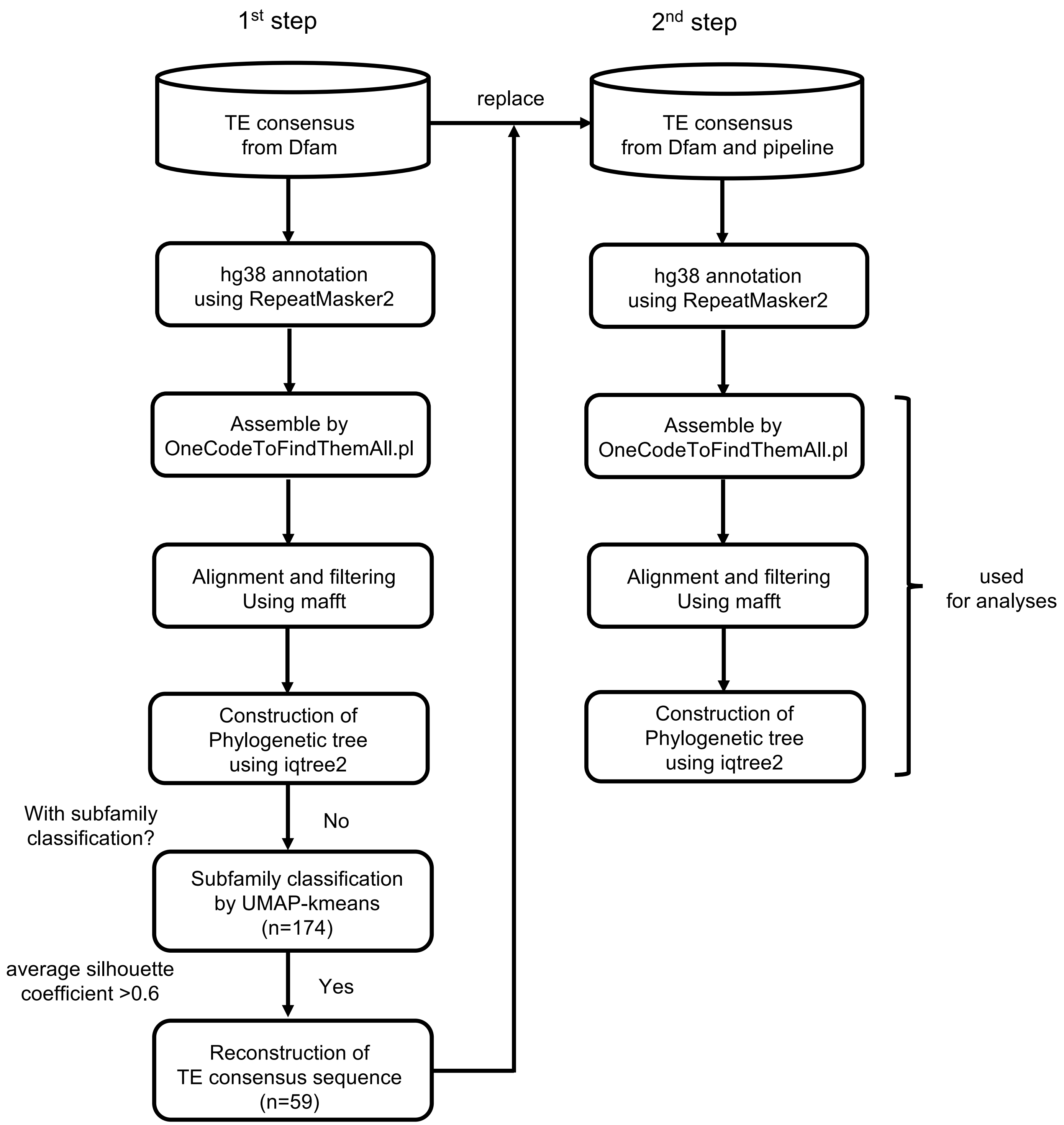


**Supplementary Fig. 3. Detail of the subfamily classification pipeline and genome annotation.** Schematic of the pipeline for subfamily classification of TE families (1st step) and genome annotation (2nd step). In the 1$^{st}$ step, we annotated the hg38 genome using consensus sequences obtained from Dfam. Subsequently, repeats were extracted from the hg38 genome, filtered, and aligned. Using the alignment data, we constructed phylogenetic trees and calculated the maximum likelihood distance matrix using the iqtree2. We then performed dimension reduction on the distance matrix using UMAP and k-means clustering. Finally, we constructed new subfamily classifications and consensus sequences for 59 TE families that met these criteria. Original consensus sequences were replaced with consensus sequences. In the 2$^{nd}$ step, a new set of consensus sequences was used to annotate the hg38 genome, which was then used for subsequent analyses. Please refer to the Methods section for further details.

Supplementary Figure 4

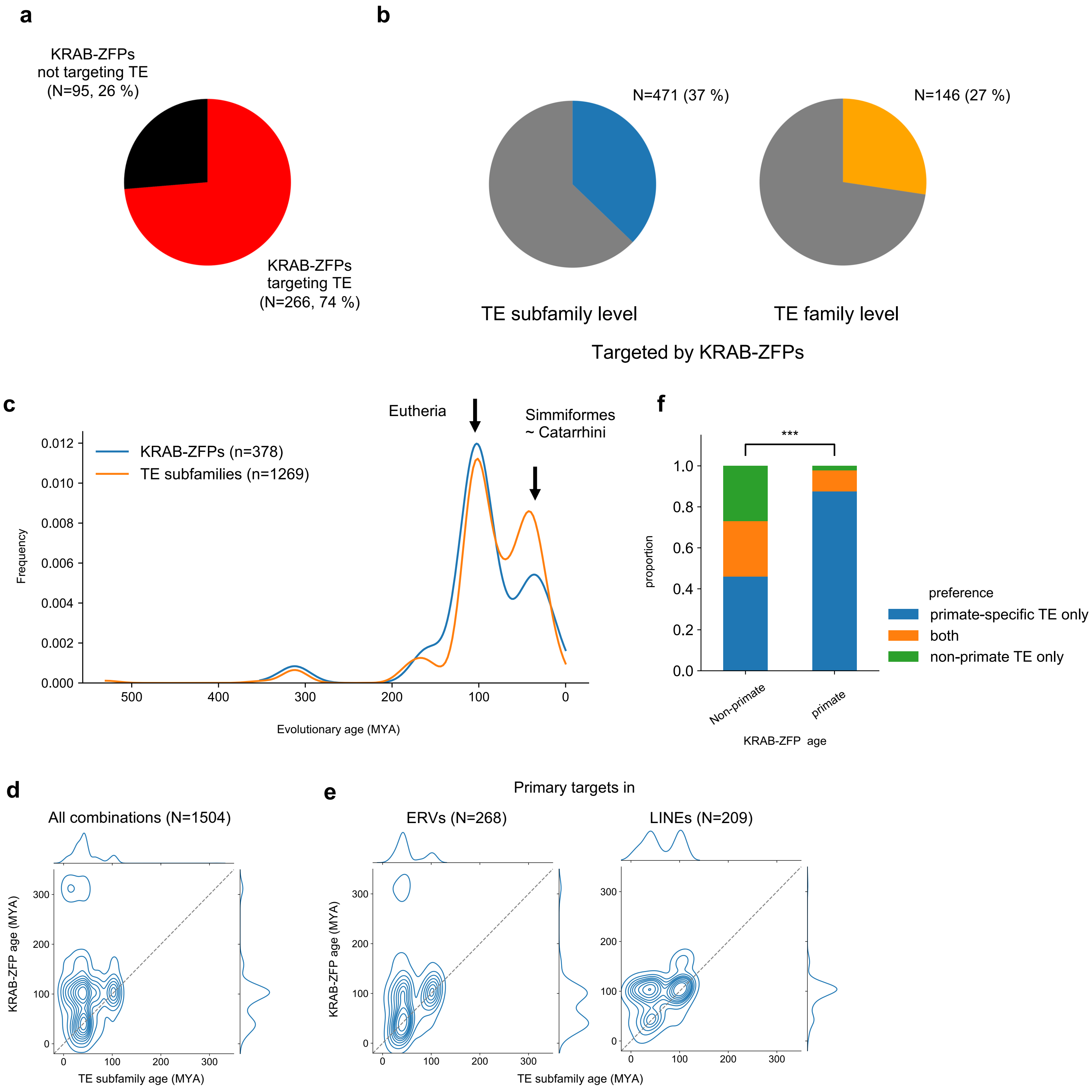


**Supplementary Fig. 4. Chronological relationship between TE families and KRAB-ZFPs.**
**a,** Proportion of KRAB-ZFP targeting TEs.
**b,** Proportion of the TE subfamily (left) and TE family (right) targeted by KRAB-ZFPs.
**c,** Evolutionary ages of KRAB-ZFPs (blue) and TE subfamilies(orange). The x-axis indicates the evolutionary age (MYA) and the y-axis indicates the probability density estimated by kernel density estimation.
**d,** Comparison of evolutionary ages between the TE subfamily (x-axis) and KRAB-ZFPs (y-axis) in 1804 primary and secondary KRAB-ZFP-TE subfamily associations.
**e,** Comparison of evolutionary ages of ERVs (left) and LINEs (right) with KRAB-ZFPs. Contour lines were estimated from 268 and 208 primary KRAB-ZFP-TE subfamily associations in ERVs and LINEs, respectively.
**f,** Preference for non-primate (left) and primate (right) KRAB-ZFPs for the evolutionary ages of the target TE subfamilies. Statistical tests were conducted using the chi-square test. ***p<0.001.

Supplementary Figure 5

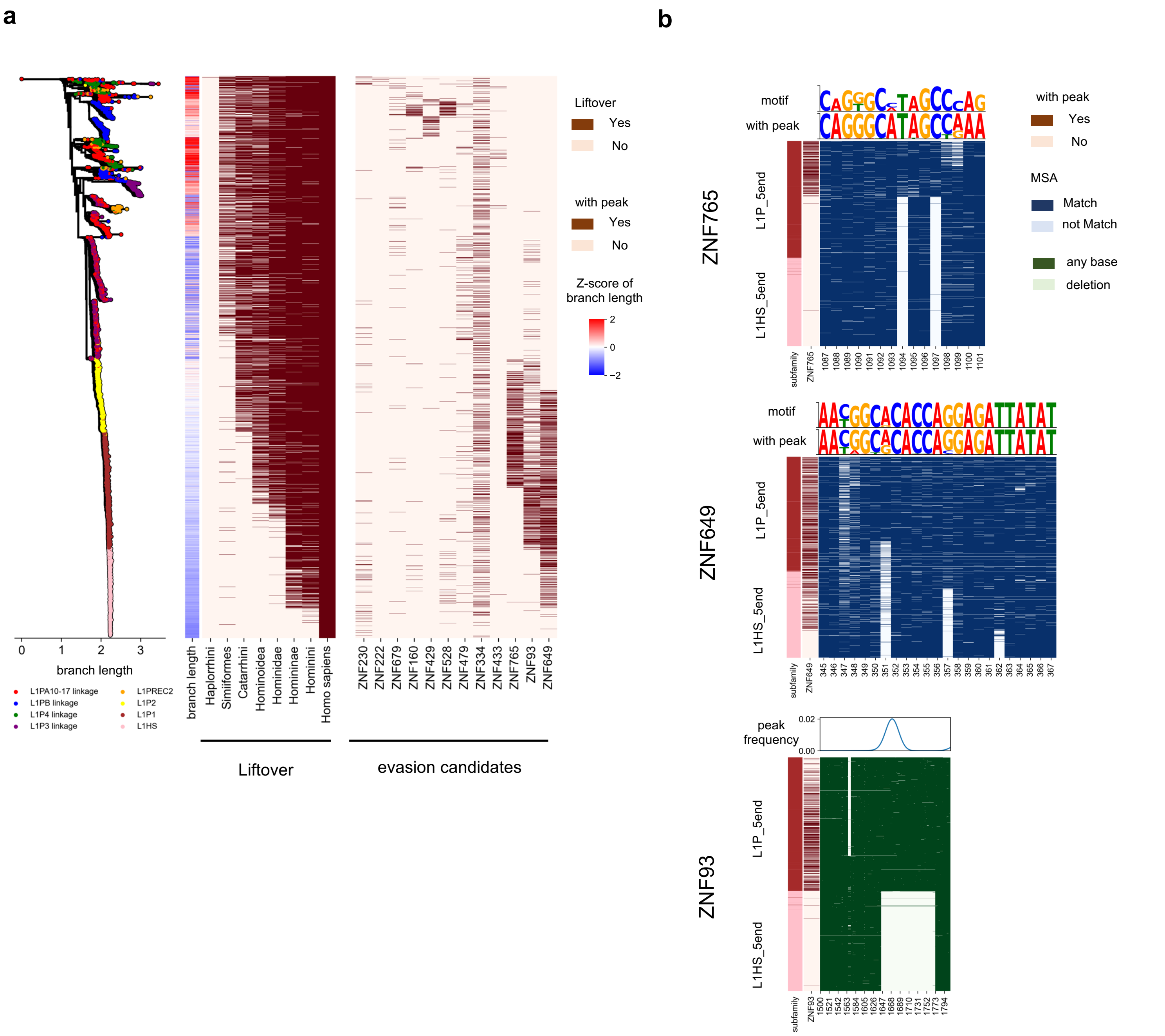


**Supplementary Fig. 5. Evasions of L1P family from KRAB-ZFPs**

**a,** Evasion of L1P_5end from KRAB-ZFPs. The phylogenetic tree indicated a phyletic relationship between the L1P_5end copies. Heatmap plots of branch length and liftover indicate the insertion date of each L1P_5end copy. The heatmap plot on the right shows the binding profile of KRAB-ZFPs that may have been evaded by L1P_5end family.

**b,** Evasion of the L1P family from ZNF765, ZNF649, and ZNF93 through point mutations and deletions. Color bars indicate the L1P_5ned subfamilies. The heatmap on the left indicates the overlap of KRAB-ZFP peaks with each L1P_5end copy. The sequence logs of ZNF765 and ZNF649 represent the de novo motif of KRAB-ZFP (upper panel) and the consensus sequence derived from L1P_5end copies with KRAB-ZFP peaks (lower panel), respectively. The ZNF93 plot shows the distribution of ZNF93 peaks. The heatmap on the right represents a match with the consensus sequence or deletion in each position and copy.

Supplementary Figure 6

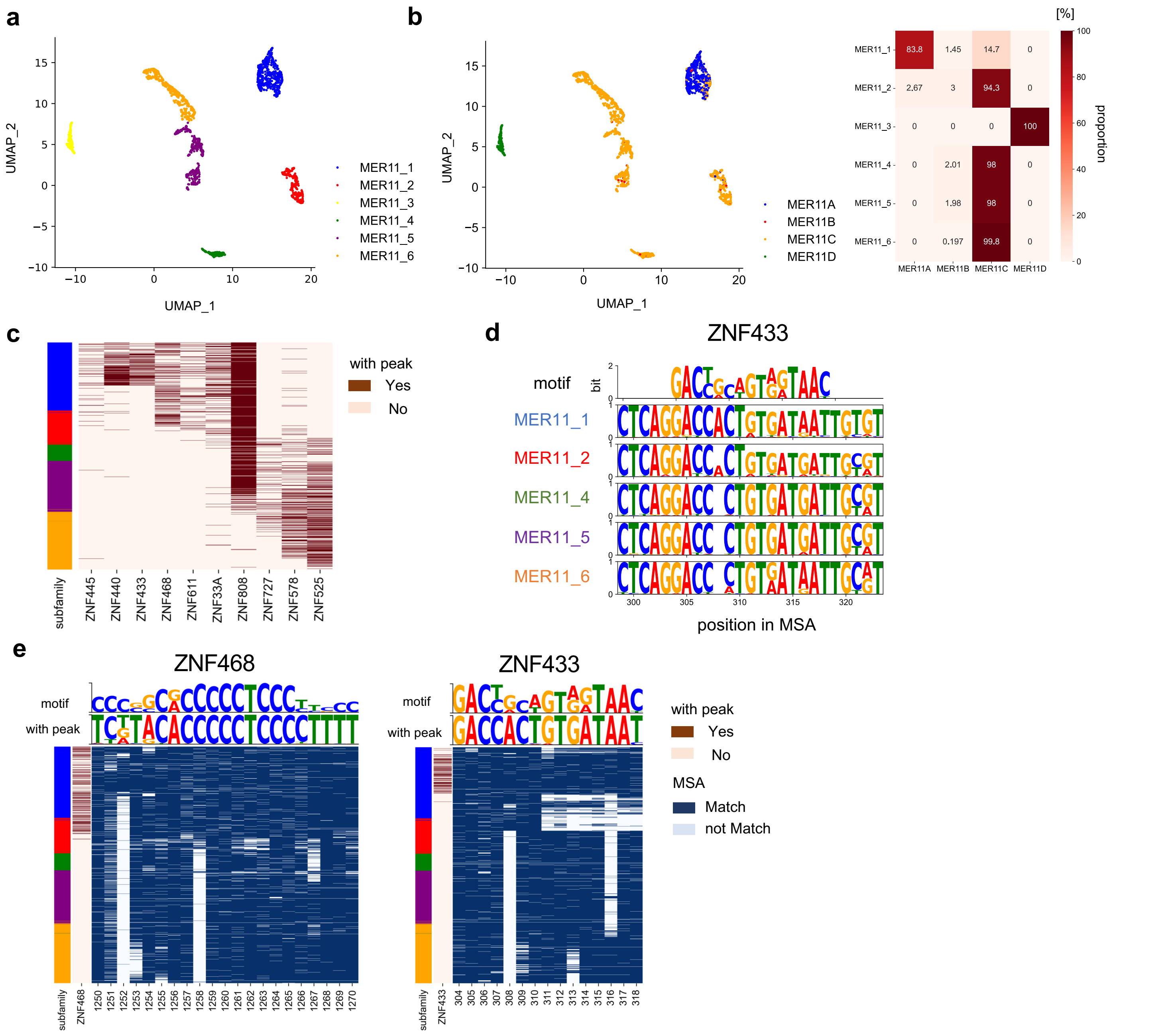


**Supplementary Fig. 6. Subfamily classification of MER11 and the evasion from KRAB-ZFPs via point mutations and deletions**

**a,** Subfamily classification of the MER11 family. The plot shows the latent space of the MER11 family obtained through dimensionality reduction using UMAP. Dots and colors represent copies and subfamilies, respectively.

**b,** Relationship between known subfamilies and subfamily classification using our pipeline. The color of each dot on the left represents a known subfamily. The heatmap and color bar represent the proportion of known subfamilies within each subfamily as determined by our pipeline. The heatmap represents the overlap between known subfamilies and each subfamily as determined by our pipeline.

**c,** Binding profile of KRAB-ZFPs to MER11 family. Color bar on the left represents the MER11 linkage in each copy. The heatmap shows the overlap of KRAB-ZFP peaks with each MER11 copy.

**d,** Evasion from ZNF433 via mutations. Sequence logos indicate the de novo motif of ZNF433 (top) and the sequence of ZNF468 binding site in the MER11 subfamily. Letters indicate the proportion of bases in each position.

**e,** Evasion from ZNF468 and ZNF433. Color bars indicate the MER11 subfamilies. The heat map on the left indicates the overlap of KRAB-ZFP peaks with each copy. The sequence logs at the top represent the de novo motif of KRAB-ZFP and the consensus sequence derived from MER11 copies with KRAB-ZFP peaks. The heatmap on the right shows a match between the consensus sequence in each position and the copy.

Supplementary Figure 7

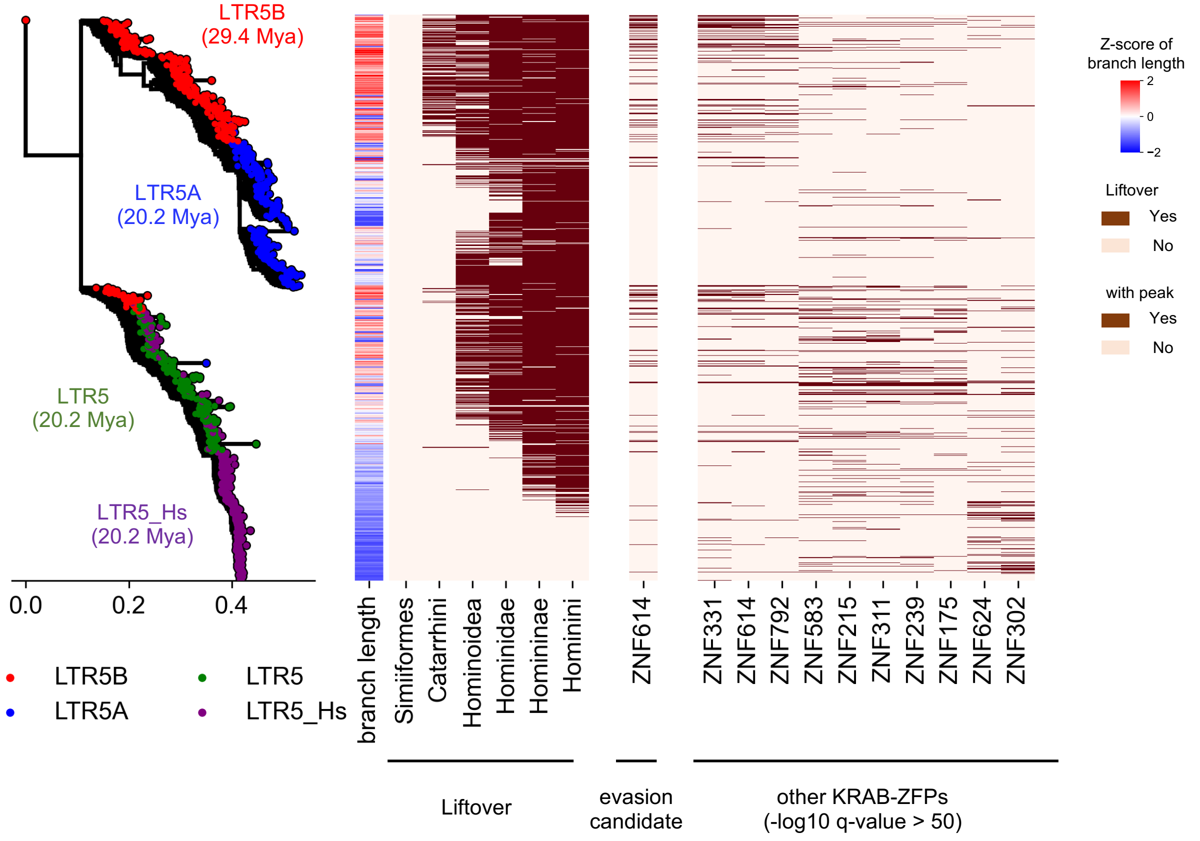


**Supplementary Fig. 7. Evolutionary arms race between LTR5 and KRAB-ZFPs.**
The phylogenetic tree indicates a phyletic relationship between the LTR5 copies. Heatmap plots of branch lengths and liftovers indicate the insertion date of each LTR5 copy. The heatmap plot on the right shows the binding profile of KRAB-ZFPs that may have evaded or targeted LTR5 subfamilies with -log10 FDR > 50.

Supplementary Figure 8

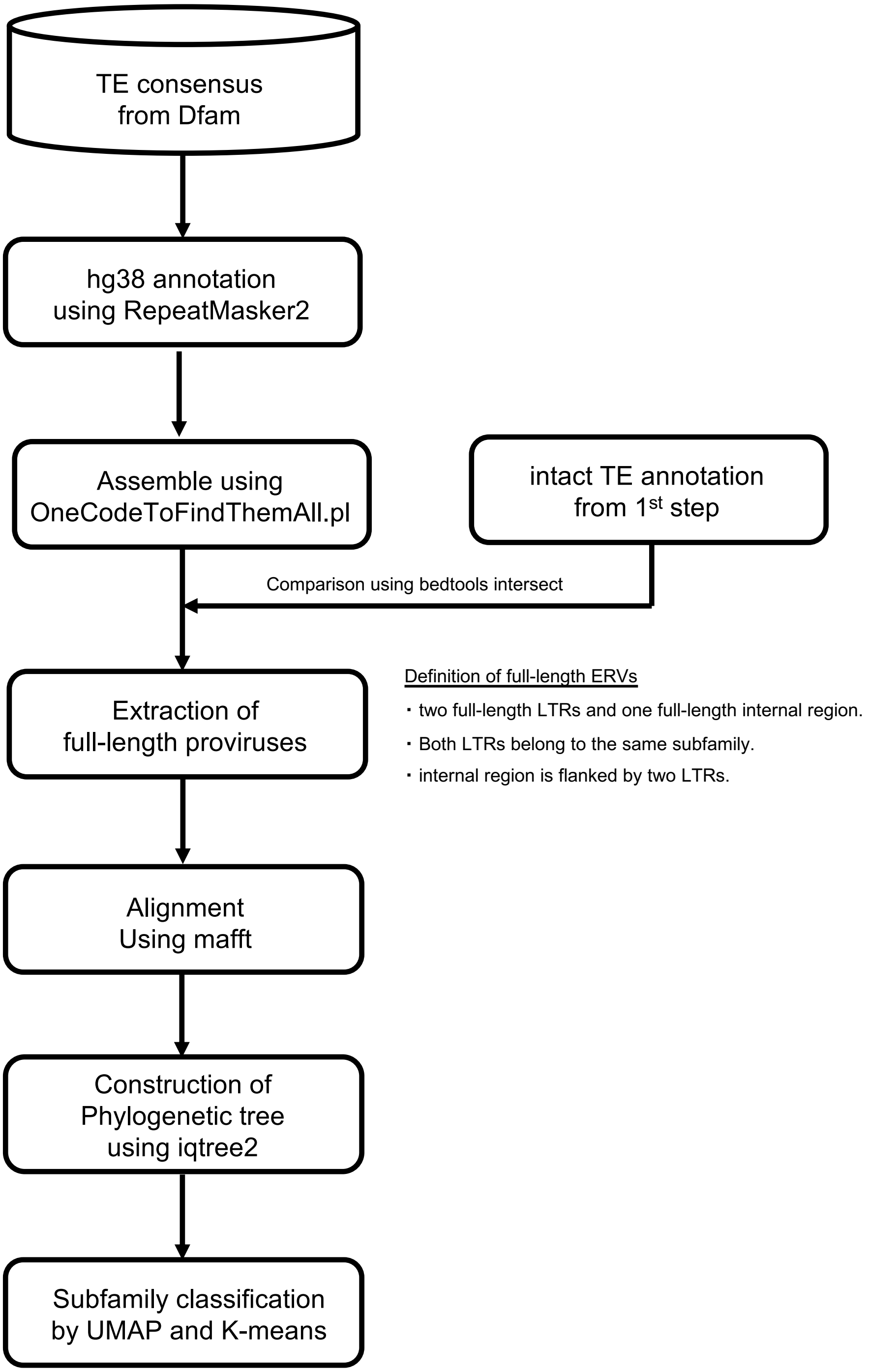


**Supplementary Fig. 8. Detail of the pipeline for whole genome provirus annotation**
Schematic of the pipeline for the identification of proviruses and subfamily classifications. The LTRs and internal regions were assembled using OneCodeToFindThemAll.pl. Proviruses were extracted by comparing with the TE annotations obtained in the 1st step (See Supplementary Fig.3 and Methods). For each ERV family, an MSA and phylogenetic tree of proviruses were constructed. Subsequently, subfamily classification was performed using UMAP and K-means.

Supplementary Figure 9

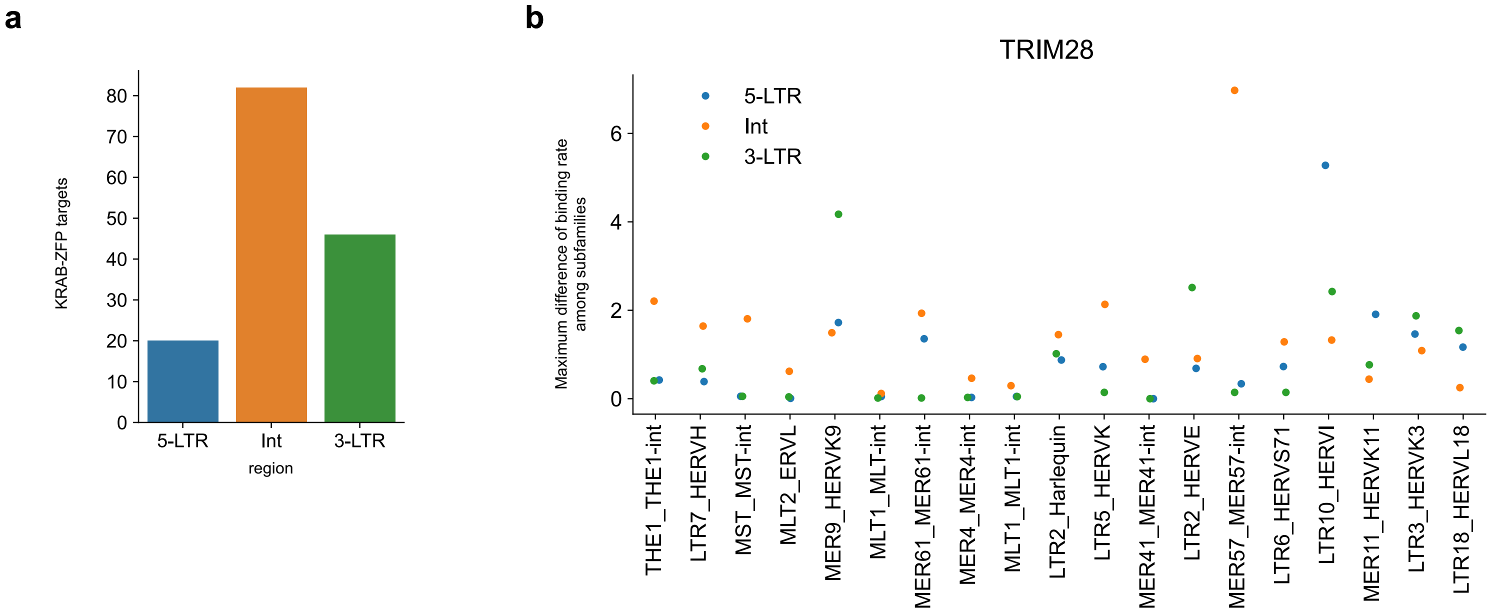


**Supplementary Fig. 9. Difference of KRAB-ZFPs/TRIM28 binding in LTRs and Internal regions.**
**a,** Number of KRAB-ZFP targets in LTRs and internal regions of full-length ERVs.
**b,** Maximum difference in TRIM28 binding rates among the subfamilies. The colors of the dots indicate the regions, including the 5-LTR (blue), internal region (orange), and 3-LTR (green).

### Supplementary Figure 10

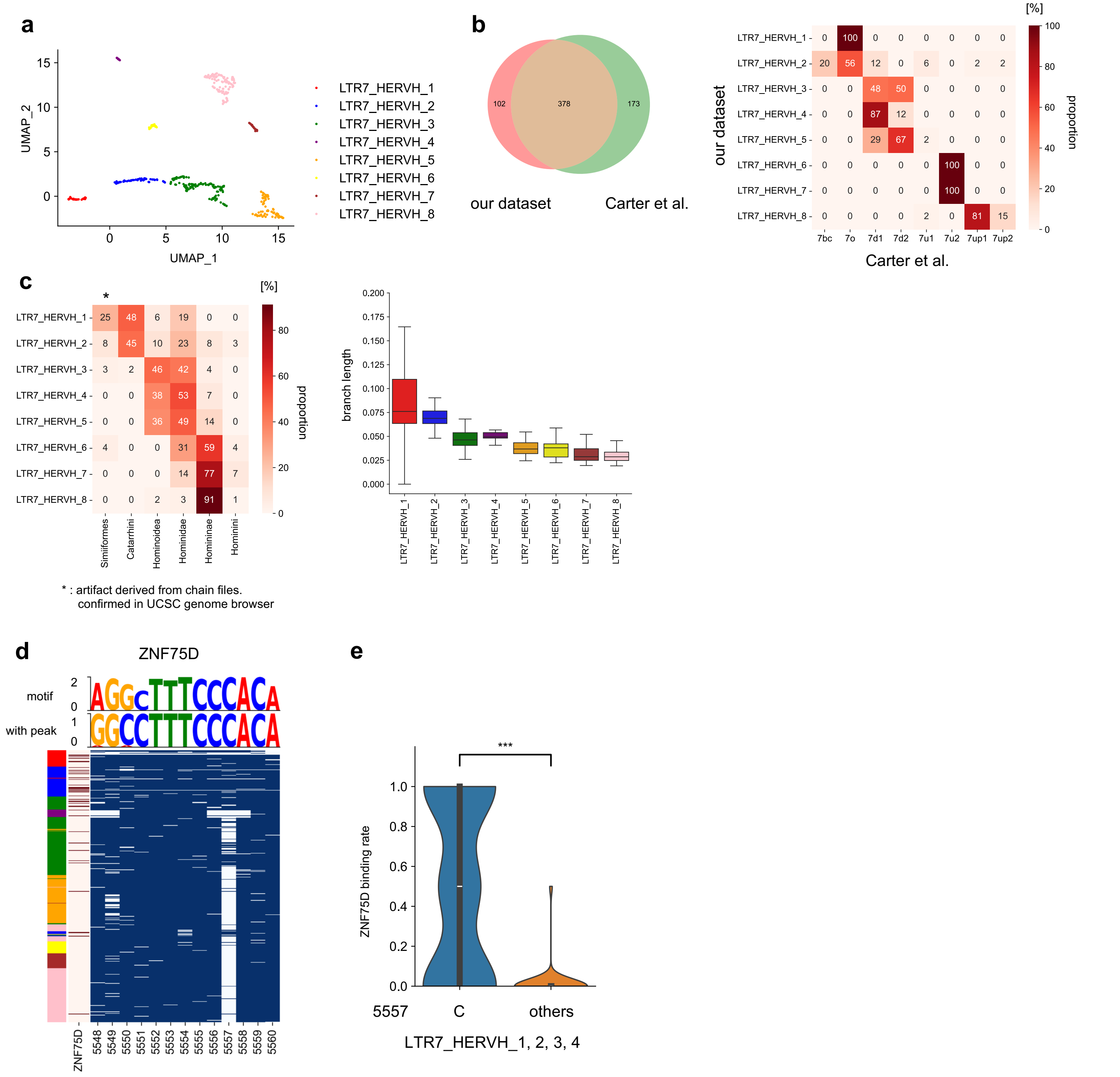


**Supplementary Fig. 10. Subfamily classification of LTR7_HERVH family and the evasion from KRAB-ZFPs.**

**a,** Subfamily classification of the LTR7_HERVH family. The plot shows the latent space of the LTR7_HERVH family obtained through dimensionality reduction using UMAP. Dots and their colors represent the copies and their respective subfamilies.

**b,** Comparison of our dataset with that of Carter et al. The left Venn diagram shows the overlap between the two datasets. Heatmap and color bar show the proportion of LTR7 subfamilies identified by Carter et al. within each subfamily, as determined by our pipeline.

**c,** Insertion date of the LTR7_HERVH subfamily. The heat map on the left shows the proportion of LTR7_HERVH copies that emerged in each era. The box plot on the right shows the branch lengths of each LTR7_HERVH subfamily.

**d,** Evasion from ZNF75D. The color bar indicates the LTR7_HERVH subfamily. The heatmap on the left indicates the overlap of ZNF75D peaks with each copy. The sequence logs at the top represent the de novo motif of ZNF75D and the consensus sequence derived from LTR7_HERVH copies with ZNF75D peaks. The heatmap on the right shows a match between the consensus sequence in each position and the copy.

**e,** Effect of the mutation on ZNF75D binding to the LTR7_HERVH family. The violin plot on the right indicates the binding rate of ZNF75D to the LTR7_HERVH_1–4 subfamily with and without the 5557 mutations. Statistical testing was conducted using the two-sided Mann–Whitney U test. ***$p<0.001$.

### Supplementary Figure 11

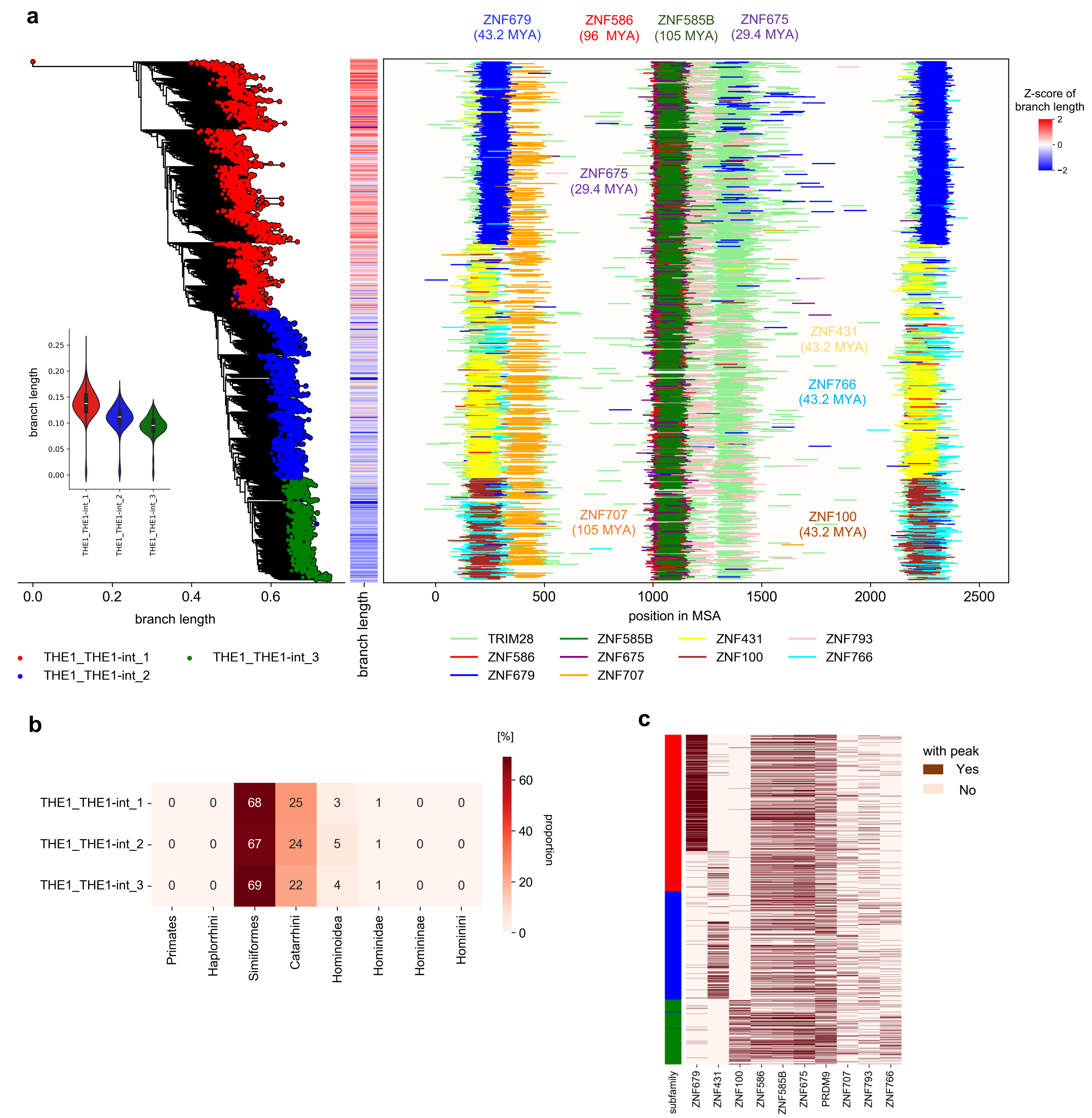


**Supplementary Fig. 11. Evolutionary arms race between THE1_THE1-int and KRAB-ZFPs.**

**a,** Evolutionary arms race between THE1_THE1-int and KRAB-ZFPs. The phylogenetic tree indicates a phyletic relationship between the THE1_THE1-int copies. Heatmap plots of the branch length and the plot on the lower left indicate the insertion date of each copy. The plot on the right shows ChIP-seq peaks of TRIM28 and KRAB-ZFPs. PRDM9 was removed because it did not have a consistent binding site.

**b,** Comparison of evolutionary ages between THE1_THE1-int subfamilies. The heat map shows the proportion of THE1_THE1-int copies that emerged in each era.

**c,** Binding profile of KRAB-ZFPs in the THE1_THE1-int family. The color bar on the left represents the THE1_THE1-int linkage in each copy. The heatmap shows the overlap of KRAB-ZFP peaks with each THE1_THE1-int copy.

Supplementary Figure 12

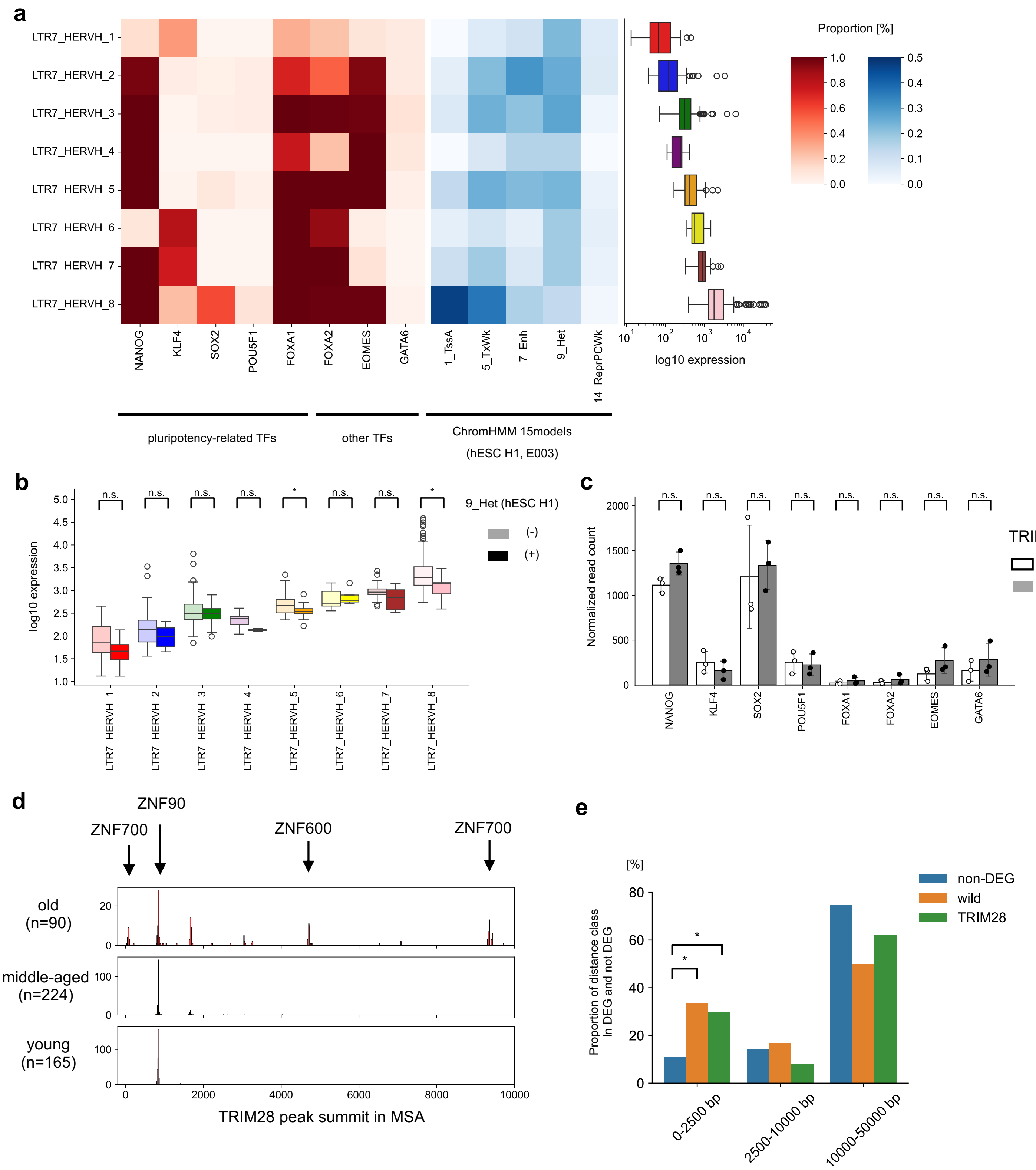


**Supplementary Fig. 12 Effect of TRIM28 deletion in hESCs on the expression of LTR7_HERVH, nearby genes, and TFs.**

**a,** Comparison of the binding patterns of TFs and chromatin state and expression in hESCs between the LTR7-HERVH subfamilies. Heatmaps show the proportion of LTR7_HERVH copies with overlapping TF peaks or chromatin states. The box plot on the right indicates the normalized read counts of LTR7-HERVH copies.

**b,** Relationship between LTR7-HERVH expression and heterochromatin state. The y-axis indicates the normalized read count of LTR7-HERVH copies. Darker and lighter colors indicate whether there was an overlap with the heterochromatin state (9_Het) in hESCs (E003). Statistical testing was conducted using the two-sided Mann-Whitney U test. P-values were adjusted using the Benjamini-Hochberg procedure. *FDR<0.05.

**c,** Effect of TRIM28 KO on TF expression. The white bars and black bars represent the means ± SD of the normalized read counts of TFs in TRIM28 wild and KO hESCs, respectively. Statistical testing was performed using DESeq2. n.s. not significant.

**d,** Histogram of TRIM28 peak summits in each LTR7_HERVH group of hESCs. The number of copies in each LTR7_HERVH group is indicated by "N=". Arrows and gene symbols of KRAB-ZFPs above the histogram indicate TRIM28 peaks related to KRAB-ZFPs.

**e,** Distance distribution of DEGs and non-DEGs within 50 kbp of LTR7_HERVH copies. The x- and y-axes represent the distance classes from the LTR7_HERVH copies and the proportion of distance classes in nearby genes, respectively. "wild" and "TRIM28" represent the genes upregulated in TRIM28 wild-type and KO hESCs, respectively. Statistical analysis was conducted using the two-sided binomial test to compare the proportions of non-DEGs. P-values were adjusted using the Benjamini-Hochberg procedure. **FDR<0.01.
